# A respiratory microdomain links ubiquinone biosynthesis to succinate dehydrogenase in an apicomplexan mitochondrion lacking Complex I

**DOI:** 10.64898/2026.10.31.763378

**Authors:** Baihetiya Baierna, Silvia NJ Moreno

## Abstract

Mitochondrial complex I (CI) is a major entry point for NADH-derived electrons and an important site of redox regulation in most eukaryotes. However, apicomplexan parasites, including *Toxoplasma gondii*, lack CI, raising a fundamental question of how mitochondrial respiration is organized in these parasites. Here, we identify a parasite-specific mitochondrial microdomain that physically and functionally couples ubiquinone (UQ) biosynthesis to succinate dehydrogenase (SDH). Using proximity labeling, membrane-based yeast two-hybrid analyses, reciprocal co-immunoprecipitations, and blue native PAGE, we show that UQ biosynthesis proteins associate with parasite-specific SDH subunits and co-migrate as a high-molecular-weight assembly. Knockdown of TgCoq4, a component of the UQ biosynthesis pathway, reduced SDH activity, and adding exogenous UQ to the assay did not restore it, indicating that UQ availability is not sufficient for efficient succinate oxidation. FCCP stimulates malate- and TMPD/ascorbate-supported respiration to maximal capacity but fails to do so with succinate as the substrate. This result, combined with reactive oxygen species (ROS) data, suggests that succinate causes over-reduction of the UQ pool, limiting further electron flow. We also find that CIII associates with the UQ synthesis machinery, suggesting a larger protein assembly. Together, these findings reveal that UQ biosynthesis and electron transport are physically and functionally integrated in *T. gondii*, defining a previously unrecognized organization of mitochondrial respiration in an organism lacking CI.

## Introduction

Mitochondria are the primary site of energy production in most eukaryotic cells, generating ATP through oxidative phosphorylation (OXPHOS). This process depends on respiratory chain complexes (RCCs) embedded in the inner membrane, which transfer electrons while pumping protons across the membrane to generate the proton motive force that drives ATP synthesis [1, 2]. Ubiquinone (UQ, coenzyme Q) occupies a central position in this process, accepting electrons from multiple dehydrogenases [3, 4] and transferring them to complex III of the respiratory chain [5]. In many organisms, RCCs assemble into higher-order supercomplexes, which have been proposed to stabilize individual complexes, enhance electron transport and reduce reactive oxygen species (ROS) production by sequestering reactive intermediates [6–8]. Although blue native PAGE (BN-PAGE) analysis fails to detect succinate dehydrogenase (SDH) in stable supercomplexes, crosslinking experiments have revealed transient or weak interactions between SDH and other ETC complexes [9].

Apicomplexan parasites are responsible for major human diseases such as malaria, cryptosporidiosis and toxoplasmosis. *Toxoplasma gondii*, the causative agent of toxoplasmosis [10], possesses a single mitochondrion that is essential for parasite replication and survival [11]. Despite having a functional ETC, *T. gondii* lacks complex I (CI)[12] and instead encodes two non-proton-pumping type II NADH dehydrogenases [13]. In addition, its mitochondrion harbors numerous essential apicomplexan-specific proteins, including components of the ETC that are absent from the host [14, 15]. Previous studies identified a supercomplex in *T. gondii*, composed of dimeric complex III (CIII_2_) and complex IV (CIV), and its assembly is important for parasite survival [16]. A separate study suggested that CII forms a supercomplex with an unknown partner, but the composition and functional significance of this complex remain unknown [17].

UQ is the central electron carrier of the ETC and serves as the first common electron carrier in OXPHOS, shuttling electrons from several dehydrogenases to CIII [18]. UQ availability and the redox state of the UQ pool (UQH_2_/UQ) are therefore central to cellular bioenergetics [4]. UQ synthesis in *T. gondii* is essential for mitochondrial function and parasite fitness, and the enzymes responsible for UQ ring modification assemble into a high-molecular-weight mitochondrial complex [19]. Notably, this complex is approximately twice the size of its mammalian counterpart, suggesting that it is part of a higher-order mitochondrial assembly with functions beyond UQ biosynthesis.

In this study, we combined TurboID proximity labeling, membrane-based yeast two-hybrid, analysis, endogenous reciprocal co-immunoprecipitation, and blue native PAGE to define how the UQ biosynthesis complex is organized in relation to the respiratory chain in *T. gondii.* We further investigated the functional significance of its association with the respiratory chain components using enzymatic assays and bioenergetic analyses. Together, this work uncovers a previously unrecognized organization of the *T. gondii* ETC that links UQ biosynthesis with succinate oxidation and shapes mitochondrial physiology in a CI-deficient parasite.

## Results

### TgCoq4 proximity labeling recovers UQ biosynthesis proteins and identifies SDH subunits

We previously identified a mitochondrial UQ synthesis complex in *T. gondii* consisting of at least five proteins (TgCoq3, TgCoq4, TgCoq5, TgCoqFMO, and TgCoq8). Although TgCoq4 was not detected in the TgCoq5-TID proximity proteome, BN-PAGE analysis showed that TgCoq4 and TgCoq5 co-migrate in a similar high-molecular complex [19]. To further investigate the composition of this complex, we fused TurboID, a promiscuous biotin ligase, to the C-terminus of TgCoq4 (TGGT1_275980) (**Fig. 1A**). We first confirmed that the *TgCoq4-TID* line biotinylates mitochondrial proteins upon incubation with biotin (**Fig. 1B**). Time-course immunofluorescence assays (IFA) further showed that *TgCoq4-TID*-mediated biotinylation specifically labels the mitochondrion after 1 h of biotin incubation (**Fig. S1**). To avoid excess labeling, we used a 1 h incubation for subsequent experiments. To identify the TgCoq4 proximity proteome, we performed subcellular fractionation to enrich for mitochondrial membranes, and to reduce cytosolic contaminants, we used a TurboID-tagged cytosolic protein as a spatial reference, as previously described [19]. After solubilizing the mitochondrial fractions, we performed streptavidin immunoprecipitation (IP) to capture biotinylated proteins (**Fig. 1C**). Mass spectrometry revealed that peptides enriched in the *TgCoq4-TID* IP relative to the cytosolic-TID control included TgCoq4, TgCoq3, TgCoq5, TgCoqFMO, and TgCoq8 (**Fig. 1D**).

**Figure 1.**
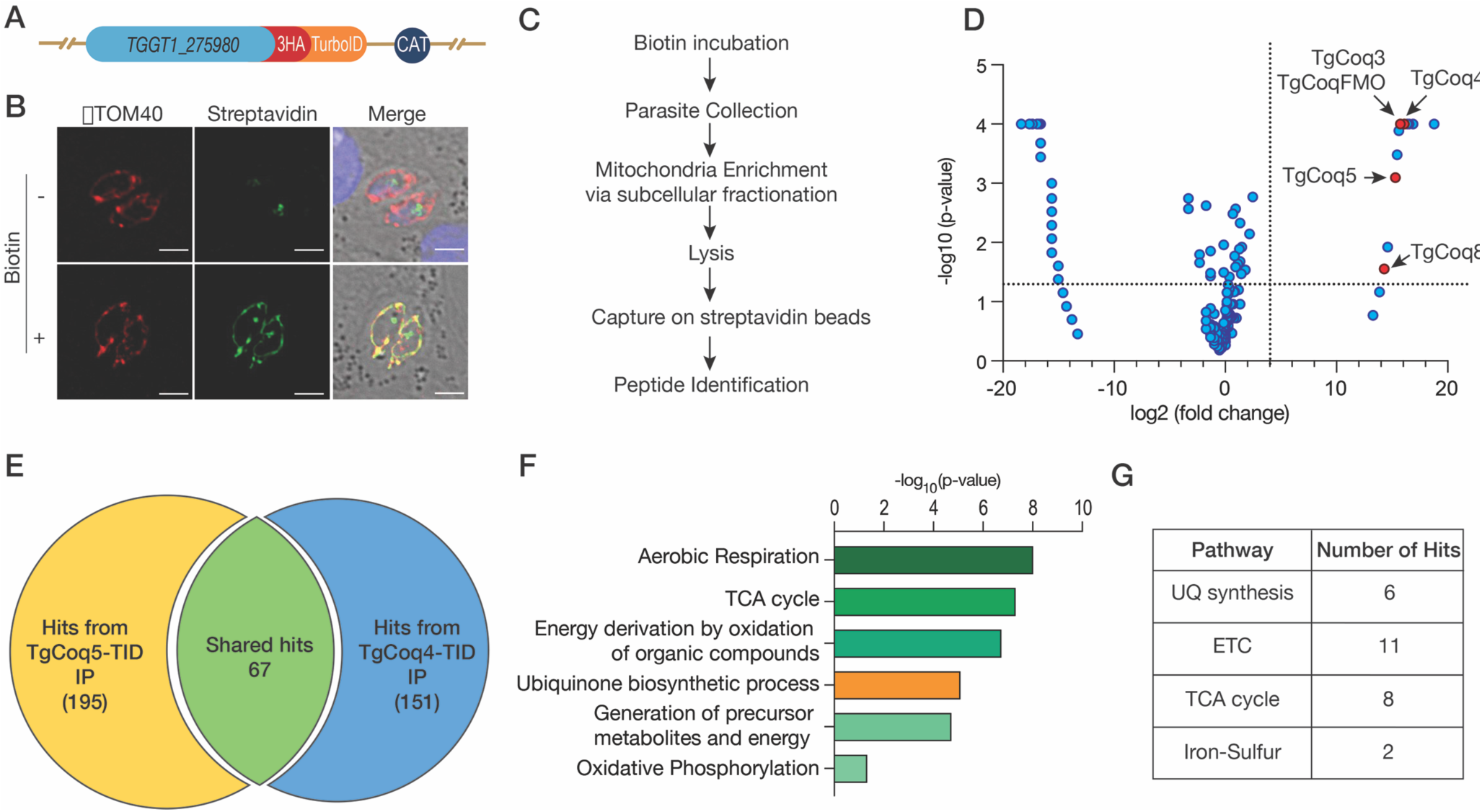
The UQ synthesis complex is in close proximity to OXPHOS proteins in *T. gondii.* (A) Schematic showing the tagging of *TgCoq4* with 3xHA-TurboID. CAT: chloramphenicol acetyl transferase gene used for selection. (B) IFAs of the *TgCoq4-TID* line showing mitochondrial protein biotinylation after 1 h of biotin incubation. TOM40 is used as mitochondrial marker, and streptavidin detects biotinylated proteins. Scale bars represent 5 μm. (C) Diagram showing the workflow used to identify the TgCoq4 proximity proteome. (D) Volcano plot of the proteins identified by mass spectrometry from the streptavidin pull-down, plotted by fold change (TgCoq4-TID/FbxO14-TID) and *p* -values. Red dots indicate UQ synthesis proteins significantly enriched at least 2-fold in *TgCoq4-TID*. (E) Venn diagram showing the overlap between proteins identified in the *TgCoq5-TID* and *TgCoq4-TID* proximity proteomes. (F) GO term enrichment analysis of the shared proteins (G) Number of proteins

Since we detected TgCoq5 in the *TgCoq4-TID* IP, we re-analyzed the *TgCoq5-TID* mass spec dataset and by lowering the minimum peptide detection threshold from two peptides to one, identified TgCoq4 peptides enriched in the *TgCoq5-TID* IP. Comparison of the two datasets revealed 67 shared hits between the TgCoq5 and TgCoq4 proximity proteomes. GO term enrichment analysis indicated that these shared proteins were enriched for terms related to aerobic respiration, tricarboxylic acid cycle (TCA), UQ biosynthesis, and oxidative phosphorylation (**Fig. 1E and 1F**). To further define their functional distribution, we classified the shared proteins according to their role in mitochondrial processes, and identified six proteins involved in UQ synthesis, 11 in the ETC, eight in the TCA cycle, and two in iron-sulfur cluster synthesis (**Fig. 1G**). Notably, most shared proteins participate in OXPHOS, suggesting that the OXPHOS machinery may form a highly interconnected network in *T. gondii*. Most strikingly, multiple SDH subunits were enriched in both proteomes, suggesting that the UQ complex could interact with SDH in *T. gondii*.

### Components of UQ synthesis complex physically associate with succinate dehydrogenase

Previous work showed that the *T. gondii* SDH complex possesses two conserved subunits, SDHA and SDHB, plus several parasite-specific subunits replacing the canonical SDHC and SDHD [17, 20]. To investigate potential interactions between the SDH and UQ synthesis proteins, we tested protein-protein interactions between subunits of these two complexes using the split-ubiquitin membrane-based yeast two-hybrid (MbY2H) assay [21]. The split-ubiquitin system allows detection of direct physical interactions *in vivo* between membrane proteins or between membrane and soluble proteins. We used the SDH subunits (SDH10, SDH23, and SDH31) and the Rieske subunit of CIII as bait proteins. We removed the mitochondrial targeting sequence (MTS) and fused each bait to the C-terminal half of ubiquitin (Cub) followed by a transcription factor (TF), with the Cub-TF portion oriented toward the yeast cytosol. The prey proteins (lacking the MTS) were TgCoq4 or TgCoq5, fused to the N-terminal half of ubiquitin. When bait and prey proteins interact, the C- and N-terminal halves of ubiquitin are brought into close proximity, reconstituting ubiquitin, which is recognized by ubiquitin-specific proteases (UBPs). The UBPs then cleave ubiquitin and release the transcription factors, which are translocated into the nucleus to activate reporter genes (*HIS3*, *ADE2*, and *LacZ*). Expression of the reporter genes allows yeast to grow on selective media lacking histidine and adenine. The *lacZ* reporter gene encodes for beta-galactosidase (β-Gal), which can be detected in colorimetric assays [22]. We validated that the bait proteins are correctly localized to the yeast plasma membrane using VP16 antibody, and the prey proteins are expressed in the yeast cytosol using HA antibody (**Fig. S2**). SDH23 continued to autoactivate despite titration with 3-amino-1,2,4-triazole (3AT), so we excluded it from the interaction studies. SDH10, SDH31, and Rieske showed robust and specific interactions with TgCoq4 and TgCoq5 (**Fig. 2**). Together, these results suggest that TgCoq4 and TgCoq5 can physically interact with parasite-specific subunits of SDH.

**Figure 2.**
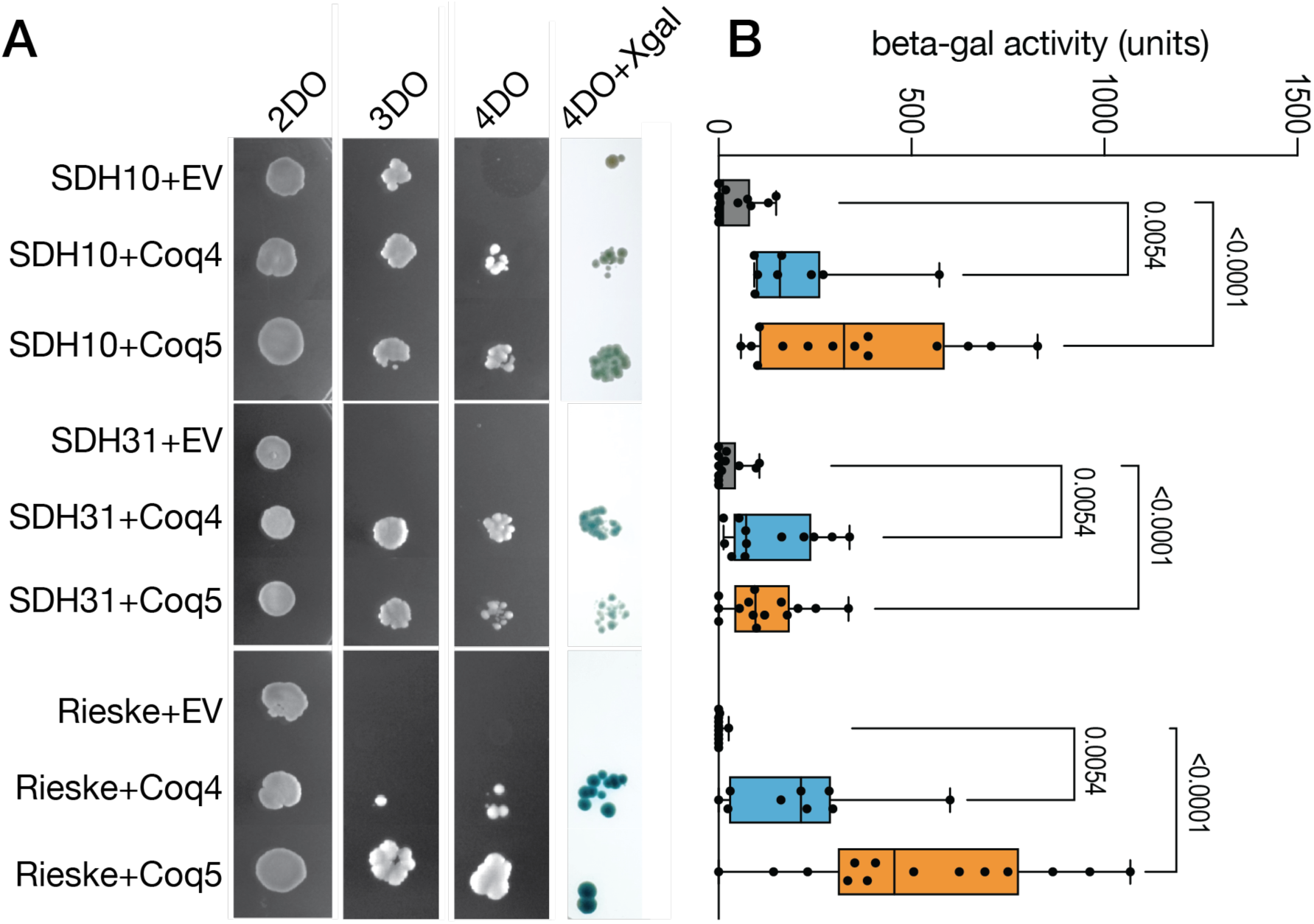
MbY2H reveals physical interactions between TgCoq4 and TgCoq5 with SDH subunits and Rieske. (A) Growth of the yeast reporter strain NMY51 expressing bait constructs (SDH10, SDH31, and Rieske), together with each prey (EV: empty vector, TgCoq4, and TgCoq5) on SD selective agar plates (SD-2DO, SD-3DO, SD-4DO, and SD-4DO+ X-Gal). A 2 μL aliquot of a 0.2 OD_600_ cell suspension was spotted on the plates and incubated at 30°C for 3 days. The experiment was repeated three times. (B) Quantitative β-galactosidase activity of the co-transformants. Data were obtained from 9-14 individual co-transformants (box and whisker plots, with center lines representing the median, and the box boundaries representing the 25^th^ and 75^th^ percentiles. Each dot represents a single co-transformant. Data were obtained from three independent experiments, and a two-way ANOVA was used for statistical analysis. For the same bait, each prey was compared to the empty vector control, and only *p*-values less than 0.05 are displayed on the figure.

### UQ synthesis complex interacts with succinate dehydrogenase and forms a supercomplex in *T. gondii*

To provide direct evidence for the interaction between the UQ synthesis complex and SDH in *T. gondii*, we performed reciprocal co-IP. We constructed parasite lines where either SDHB or SDH23 was endogenously tagged with Myc in the background of TgCoq4 or TgCoq5 endogenously tagged with HA (**Fig. 3A-B**). We then performed IPs with αHA magnetic beads to pull down either TgCoq4 or TgCoq5. In the elution, we recovered TgCoq4 or TgCoq5 and co-immunoprecipitated SDHB or SDH23 (respectively). SAG1, which was used as a negative IP control, was only present in the input and was not detected in the elution. Reciprocal IPs using αMyc magnetic beads to immunoprecipitate SDHB or SDH23, showed recovery of TgCoq4 or TgCoq5 in the elution while SAG1 was only detected in the input (**Fig. 3C-D**). These results indicate that TgCoq4 and TgCoq5 interact with SDHB and SDH23 and are likely present in the same protein complex.

**Figure 3.**
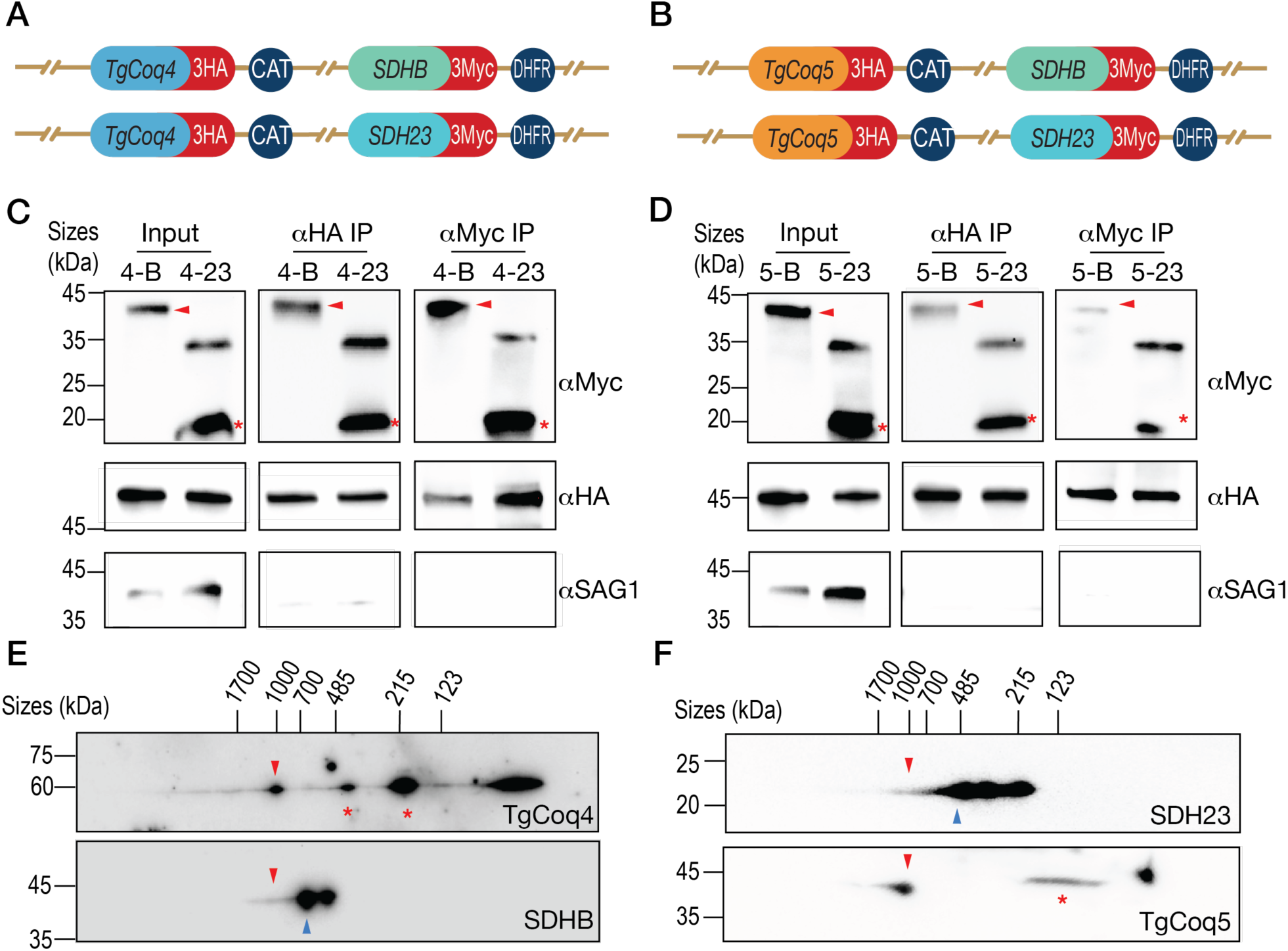
UQ synthesis complex proteins interact with SDH subunits in *T. gondii*. (A) Schematic showing the double tagging of *TgCoq4-3HA* with *SDHB-3Myc* (4-B) and *TgCoq4-3HA* with *SDH23-3Myc* (4-23). CAT: chloramphenicol acetyl transferase gene used for selection; DHFR: dihydrofolate reductase gene used for pyrimethamine resistance. (B) Schematic showing the double tagging of *TgCoq5-3HA* with *SDHB-3Myc* (5-B) and *TgCoq5-3HA* with *SDH23-3Myc* (5-23). (C) Reciprocal co-IPs using the *TgCoq4-3HA* line expressing *SDHB-3Myc* (4-B) or expressing *SDH23-3Myc* (4-23). aHA was used to pull down TgCoq4-3HA, and aMyc was used to pull down SDHB-3Myc or SDH23-3Myc. aSAG1 was used as a negative IP control. (D) Reciprocal co-IPs similar to **C**, except that the *TgCoq5-3HA* line expressing *SDHB-3Myc* (5-B) or *SDH23-3Myc* (5-23) was used. aHA was used to pull down TgCoq5-3HA, and aMyc was used to pull down SDHB-3Myc or SDH23-3Myc. For **C** and **D**, the red arrowhead indicates SDHB-3Myc, while the red asterisk indicates the SDH23-3Myc monomer. (E) Two-dimensional BN/SDS-PAGE using the double-tagged line 4-B. SDHB was detected using aMyc, while TgCoq4 was detected using aHA. (F) Two-dimensional BN/SDS-PAGE using the double-tagged line 5-23. SDH23 was detected using aMyc, while TgCoq5 was detected using aHA. For **E** and **F**, the blue arrowhead indicates the SDH, while the red arrowhead indicates the supercomplex containing the UQ synthesis complex and SDH. Red asterisks indicate UQ synthesis complex assembly intermediates or disassembly products. SDHB and SDH23 were probed with aMyc; TgCoq4 was probed with aHA, and TgCoq5 was probed with aTgCoq5. Experiments in panes **C-F** were repeated three times.

To further investigate the presence of a supercomplex, we performed BN-PAGE analysis using the double tagged mutants shown in Fig 3A. For each line, we performed two-dimensional BN/SDSPAGE (2DPAGE), where the native gel strip was cut, denatured and further separated on an SDSPAGE gel. By probing the membrane with αHA, we observed the UQ synthesis complex proteins present at ∼1 MDa. Using an αMyc antibody, we observed that a small portion of SDH co-migrates at ∼1 MDa, supporting the formation of a supercomplex (**Fig. 3E-F**). These observations demonstrate that a fraction of SDH and the UQ synthesis complex co-migrate as part of a large supercomplex in *T. gondii*.

Since the direct physical interaction was only observed with parasite specific SDH subunits, we hypothesized that this interaction is not present in mammalian host cells. To test this hypothesis, we performed BN-PAGE and 2DPAGE using commercial SDHB and Coq5 antibodies and HEK cell lysates. We observed a single band around ∼123 kDa using αSDHB, corresponding to the mammalian SDH. Probing the membrane with an αCoq5 antibody revealed a single band between 215 and 485 kDa, corresponding to the mammalian UQ synthesis complex (**Fig. S3A**). Two-dimensional PAGE further supported that SDH does not form a supercomplex under the condition tested (**Fig. S3B**). These findings indicate that the interaction observed in *T. gondii* is unique to apicomplexan parasites.

### Succinate-driven respiration does not reach maximal respiratory capacity

We noticed that succinate-driven respiration in *T. gondii* differs from that of organisms containing complex I. In organisms containing CI, such as humans, FCCP stimulates succinate-driven mOCR to its maximal capacity [23]. However, in *T. gondii*, FCCP fails to increase succinate-driven mOCR to its maximal capacity [12] (**Fig. 4A-B**). To verify that this phenotype was not due to incomplete uncoupling, we used safranin *O* as a proxy for mitochondrial membrane potential and found that FCCP successfully dissipated the membrane potential under the same conditions (**Fig. S4A-B**). We titrated succinate and FCCP to confirm that the observed phenotype was not due to suboptimal substrate or uncoupler concentrations (**Fig. S4C-D**).

**Figure 4.**
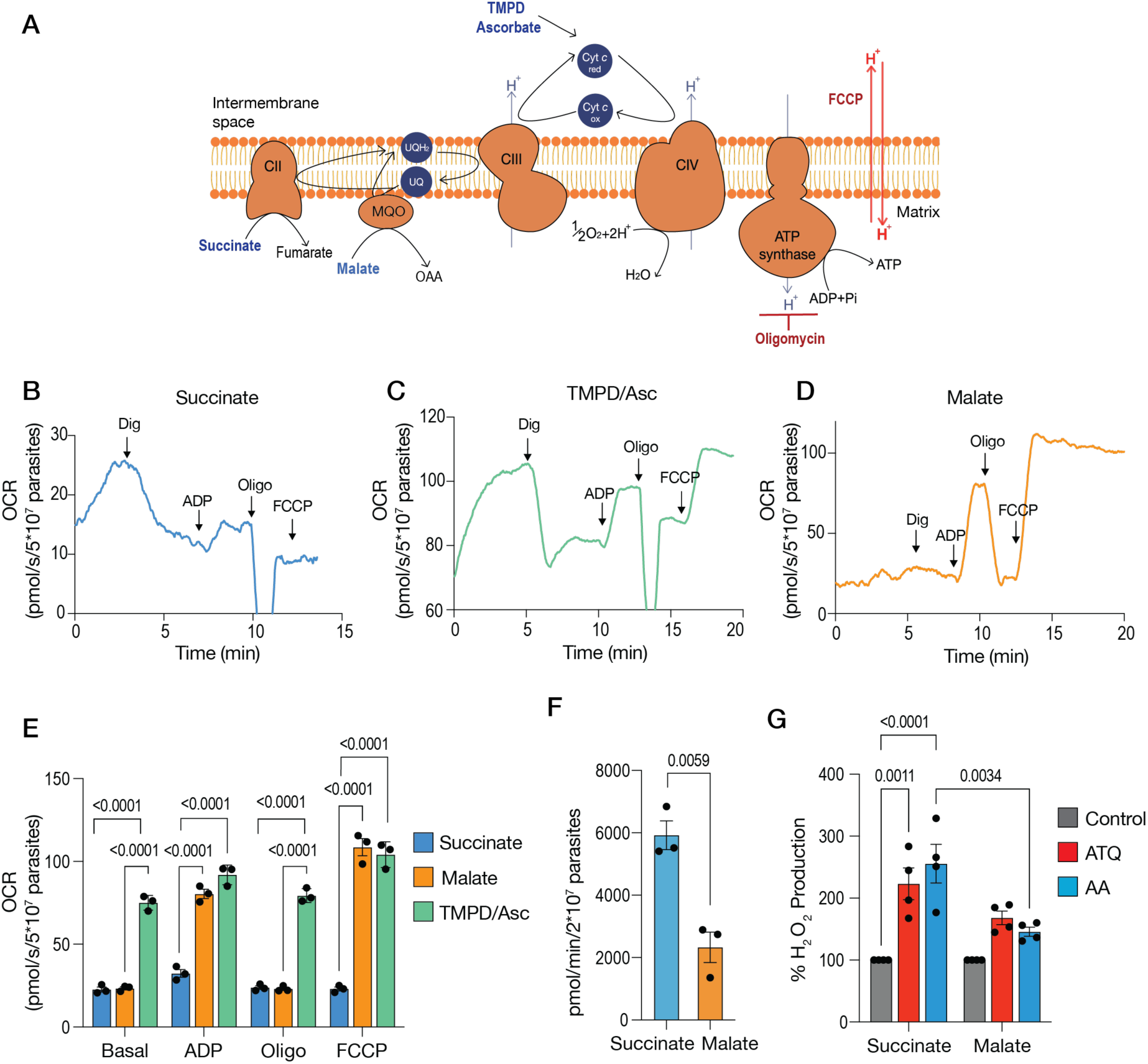
Succinate-driven mOCR fails to increase upon FCCP uncoupling and is associated with increased ROS production. (A) Schematic of the *T. gondii* ETC with substrates and inhibitors indicated. (B) Representative mOCR trace using 2 mM succinate as the substrate. Dig: digitonin, oligo: oligomycin. The concentrations used are listed in the Methods section. (C) Representative mOCR trace using 10 mM malate as the substrate. (D) Representative mOCR trace using 0.2 mM TMPD and 1.5 mM ascorbate as the substrates. (E) Quantification of basal, ADP-induced, oligomycin-inhibited and FCCP-stimulated mOCR using the substrates shown in **B-D**. (F) Cytochrome *c* reduction activity using either succinate or malate as the substrate. (G) Percentage increase in H_2_O_2_ using succinate or malate as the substrate in the absence or presence of ATQ (atovaquone) or AA (antimycin A) as inhibitors. For **E**-**F**, data are presented as mean ± SEM from three biological replicates. For **G**, data are represented as mean ± SEM from four biological replicates. A two-way ANOVA was used for **E** and **G**, while a t-test was used for **F**.

To rule out CIV as the bottleneck responsible for this phenotype, we measured mOCR using TMPD/ascorbate to reduce cytochrome, which then transfers electrons to CIV. After FCCP addition, maximal mOCR was significantly higher with TMPD/ascorbate than with succinate (**Fig. 4C** and **4E**), indicating that CIV is not the rate limiting step. Next, we tested malate, which supplies electrons to UQ through malate: quinone oxidoreductase (MQO). Both ADP and FCCP-stimulated mOCR were significantly higher with malate than with succinate (**Fig. 4D-E**, **Table 1**). This suggests that the respiratory chain can support higher mOCR when electrons enter the UQ pool through MQO, and that the limitation is specific to succinate-driven respiration.

**Table 1.**
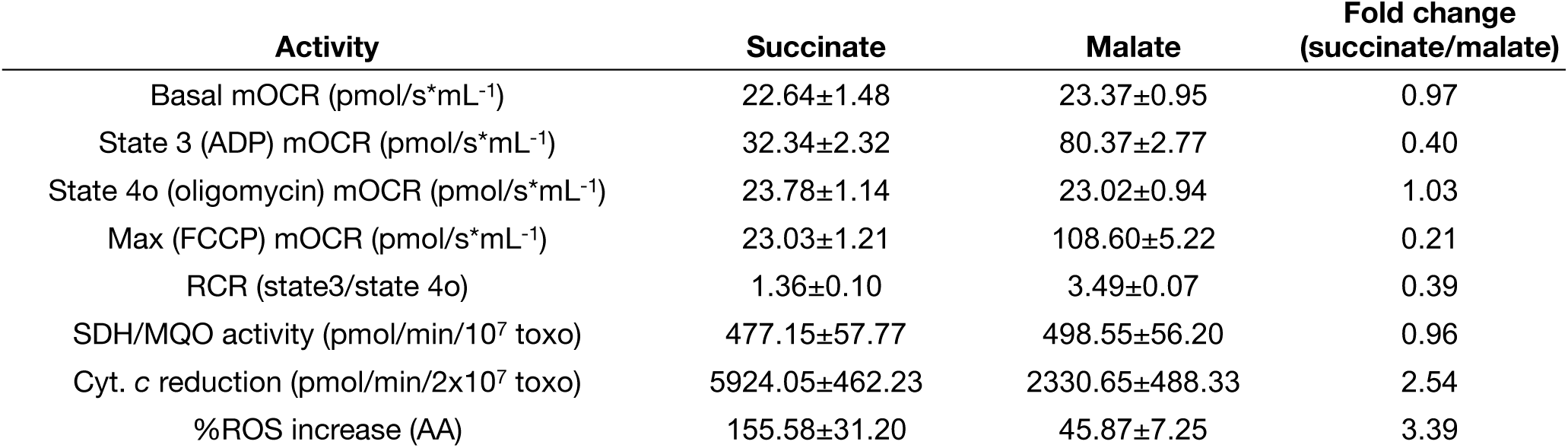
Comparison of succinate- and malate-supported mitochondrial activities in *T. gondii.* Summary table showing the different respiratory states, RCR (respiration control ratio), activities of individual respiratory complexes, cytochrome *c* reduction, and the percentage increase in ROS upon AA inhibition using either succinate or malate as the substrate. Data are represented as mean ± SEM from three biological replicates.

Intriguingly, the inability of succinate to increase respiration following FCCP addition was not due to a low initial rate of electron transfer through the succinate pathway. The initial rate of cytochrome *c* reduction was significantly higher with succinate than with malate (**Fig. 4F**, **Table 1**). This finding demonstrates that the succinate pathway supports robust initial electron transfer activity. Because succinate concentration is linked to the UQH₂/UQ ratio [24], we reasoned that the insensitivity of succinate-driven mOCR to FCCP stimulation may be caused by the UQ redox state. We hypothesized that succinate drives overreduction of UQ, resulting in the accumulation of UQH₂ and a decrease in oxidized UQ available to accept electrons from SDH. This could limit further succinate oxidation and explain why FCCP fails to increase mOCR.

To test this hypothesis, we assessed the redox state of the respiratory chain indirectly by measuring H_2_O_2_ production using specific CIII Q-site inhibitors. The Q_o_ and Q_i_ sites are the two UQ-binding sites of CIII that mediate UQH_2_ oxidation and UQ reduction, respectively, during the Q cycle [25]. We measured H_2_O_2_ production following inhibition of either the Q_i_ or Q_o_ using succinate or malate as substrates, in the presence of oligomycin to inhibit ATP synthase. When succinate was used as the substrate, inhibition of either the Q_o_ or Q_i_ site of CIII resulted in similarly robust increases in H_2_O_2_ production (**Fig. 4G**). By contrast, malate did not significantly increase H_2_O_2_ production under similar conditions (**Fig. 4G**, **Table 1**). Together with the respiratory analyses, these findings are consistent with succinate driving overreduction of the UQ pool, while malate is associated with a more oxidized UQ pool. Because Q_i_ and Q_o_ inhibitors can differentially affect ROS production depending on the redox state of the respiratory chain [26], the comparable responses observed with both inhibitors in the presence of succinate suggest that succinate generates substantial electron pressure on CIII.

Together, our data suggest that in an organism lacking canonical CI, succinate-driven electron input locally over-reduces the UQ pool, preventing mOCR from increasing even when uncoupling dissipates the membrane potential. These findings suggest that the interaction between the UQ synthesis complex and SDH may facilitate the local delivery of newly synthesized UQ to SDH, supporting succinate oxidation in the absence of CI.

### TgCoq4 knockdown impairs succinate dehydrogenase activity

To investigate the functional significance of the interaction between the UQ synthesis complex and SDH, we performed an SDH activity assay using succinate as the substrate and 2,6-dichlorophenolindophenol (DCPIP) as the terminal electron acceptor. Because knockdown of UQ complex subunits can decrease UQ synthesis, we supplemented the reaction buffer with exogenous UQ_6_ as the electron carrier. We generated a conditional knockdown line of SDHB, *iΔTgSDHB*, as a positive control for this assay. As expected, SDHB knockdown significantly reduced initial SDH activity (**Fig. 5A**). Interestingly, TgCoq4 knockdown caused a similar defect (**Fig. 5A**). To determine whether the decrease in SDH activity was due to poor parasite viability or an inability of UQ_6_ to support electron transfer, we used a malate-quinone oxidoreductase (MQO) activity assay as a control. MQO activity was unchanged following knockdown of either TgSDHB or TgCoq4, indicating that the decrease in SDH activity was not due to a general loss of mitochondrial enzyme activity or an inability of UQ_6_ to accept electrons (**Fig. 5B-C**, **Table 1**). Although TgCoq4 conditional knockdown mutants have an established growth defect [19], they remained viable at the time of the assay, as shown by their ability to form comparable plaques when ATc was removed following the 3-day preincubation (**Fig. S5A** and **S5B**, +Pre-ATc, *top row*). SDH activity was measured from the initial linear rate of DCPIP reduction. Although TgCoq4 knockdown markedly decreased this rate, DCPIP was eventually fully reduced when the reaction was allowed to continue, indicating that SDH remained active but at a reduced rate. In contrast, TgSDHB knockdown did not fully reduce DCPIP even at later time points.

**Figure 5.**
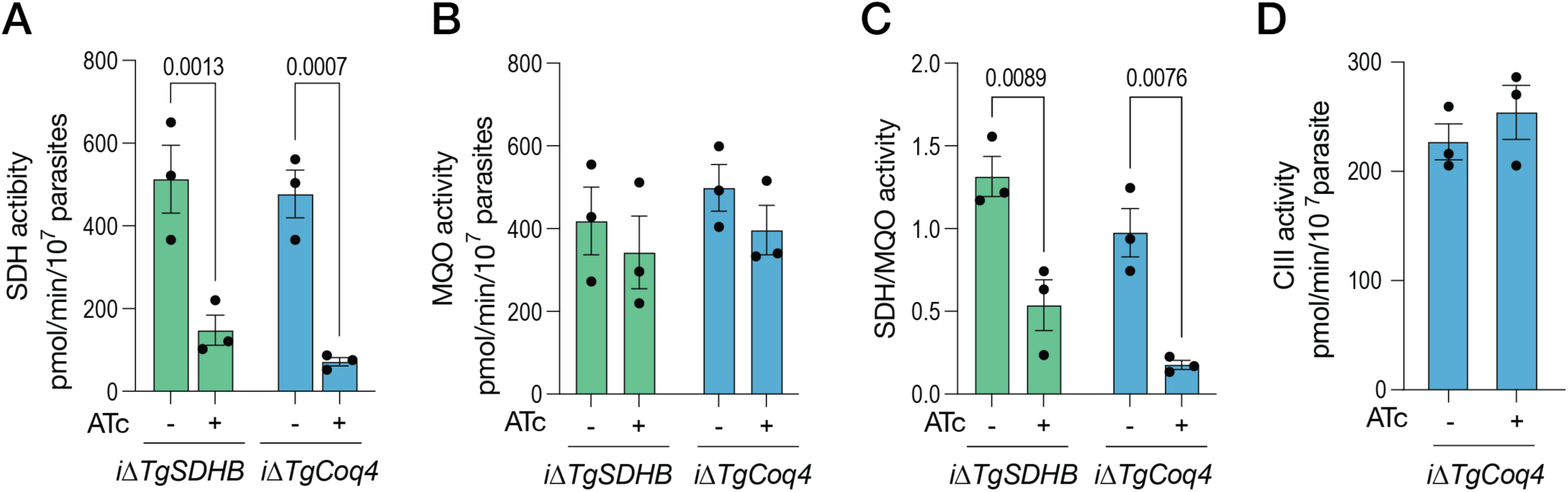
TgCoq4 is needed for optimal SDH activity. (A) Quantification of SDH activity in *iΔTgSDHB* and *iΔTgCoq4* mutants cultured without or with ATc for 3 days. The assay was performed in the presence of 10 μM UQ_6_. Data are presented as the mean ± SEM from three biological replicates. (B) Quantification of MQO activity using the same mutants and conditions as in part **A**. (C) Ratio of SDH/MQO activity using data from parts **A** and **B**. (D) CIII activity assay in *iΔTgCoq4* mutants cultured without or with ATc for 3 days. Data are presented as the mean ± SEM from three biological replicates. For A-C, a two-way ANOVA test was used, and t-test was used for D.

To investigate whether this effect was specific to TgCoq4 or shared by other UQ synthesis complex subunits, we also assayed SDH activity in the *iΔTgCoq5* line. Unlike the defect observed when TgCoq4 was knocked down, TgCoq5 knockdown did not impair SDH activity (**Fig. S5C**). We also measured CIII activity using ubiquinol as the substrate with cytochrome *c* as the electron acceptor. TgCoq4 knockdown did not cause a reduced CIII activity (**Fig. 5D**).

Contrary to the yeast UQ synthome, where knockout of a subunit impairs complex assembly [27], the *T. gondii* UQ synthesis complex remains intact upon subunit knockdown [19]. To investigate whether the impaired SDH activity results from defective SDH complex assembly or reduced SDH protein levels, we endogenously tagged SDHB with Ty in the *iΔTgCoq4* line. Interestingly, knockdown of TgCoq4 for 3 days did not impair SDH assembly or the formation of the 1 MDa supercomplex (**Fig. S6A**). Furthermore, SDHB protein levels were unaffected upon TgCoq4 knockdown (**Fig. S6B-C**), indicating that the impaired SDH activity is not due to SDH disassembly or reduced protein levels.

### TgCoq4 knockdown selectively alters substrate-dependent respiration

Since TgCoq4 knockdown selectively impaired SDH activity, we next evaluated the bioenergetics of the TgCoq4 mutant in the presence of either succinate or malate as substrate to determine if this defect affected mitochondrial function differently depending on the substrate used. As expected, knockdown of TgCoq4 significantly reduced respiration regardless of the substrate (**Fig. 6A-C**). However, the basal membrane potential, estimated from the difference between the signal before ADP addition and after FCCP addition, was significantly reduced only when succinate was the substrate. Malate-driven basal membrane potential was unchanged after TgCoq4 knockdown (**Fig. 6D-F**).

**Figure 6.**
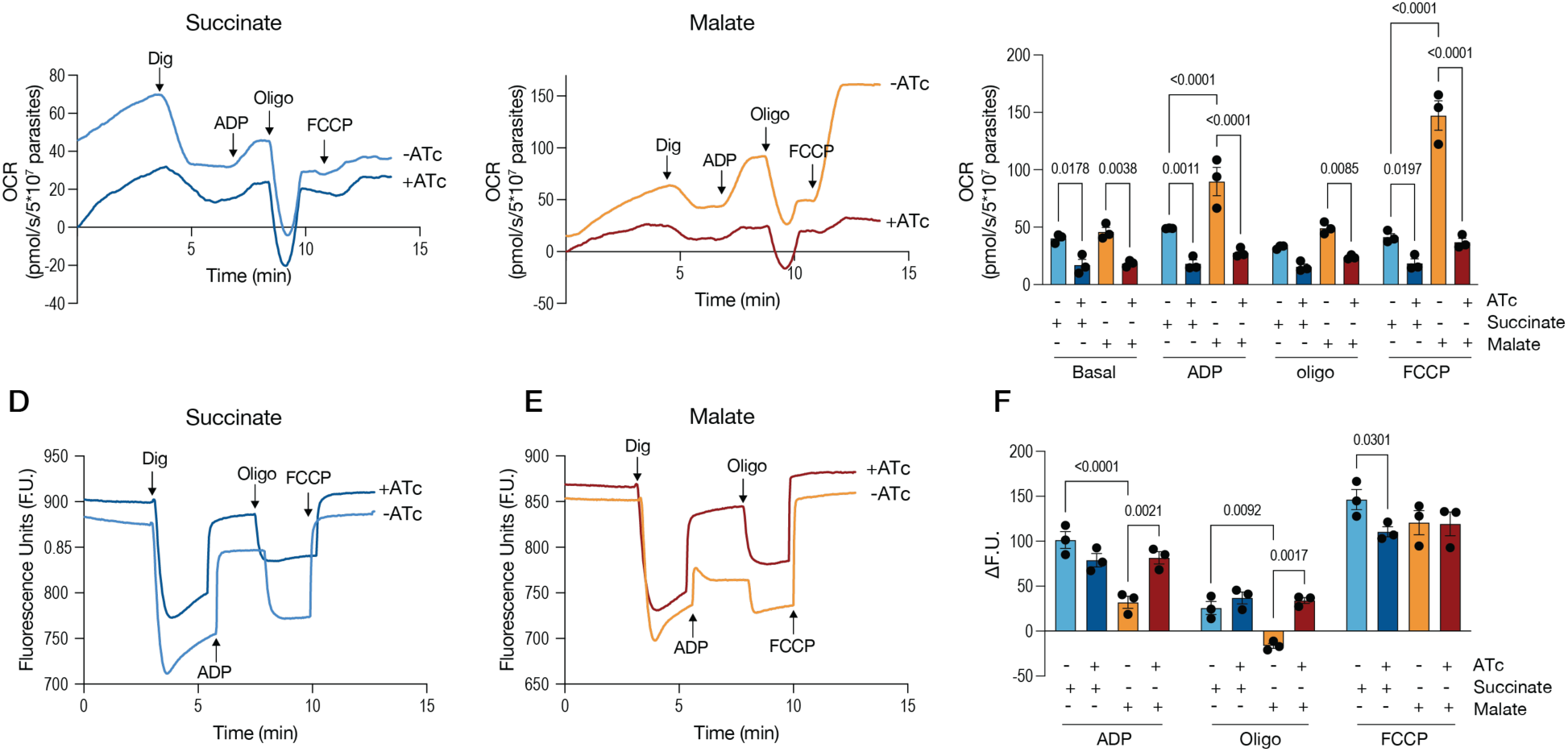
TgCoq4 knockdown selectively collapses succinate-driven membrane potential. (A) Representative mOCR traces of *iΔTgCoq4* mutants using succinate as substrate. -ATc: *iΔTgCoq4* cultured without ATc; +ATc: *iΔTgCoq4* cultured with ATc for 2 days. (B) Representative mOCR traces similar to **A**, except that malate was used as the substrate. (C) Quantification of the different mOCR states from the conditions shown in **A** and **B**. (D) Representative safranin O fluorescence traces of *iΔTgCoq4* mutants using succinate as the substrate. -ATc: *iΔTgCoq4* cultured without ATc; +ATc: *iΔTgCoq4* cultured with ATc for 2 days. (E) Representative safranin O fluorescence traces similar to **D**, except that malate was used as the substrate. (F) Quantification of fluorescence (F.U.) changes in ADP (ADP-basal), oligomycin (oligomycin-basal), and FCCP (FCCP-basal) responses from the conditions shown in **D** and **E**. For C and F, data are presented as mean ± SEM from three biological replicates. A two-way ANOVA was used for statistical analysis.

We next examined the response of the membrane potential to ADP and oligomycin to assess how TgCoq4 knockdown affects oxidative phosphorylation. Interestingly, ADP caused a smaller membrane depolarization when malate was used as the substrate than when succinate was used (**Fig. 6F**, -ATc, ADP). This suggests that malate-supported respiration is more effective at maintaining the membrane potential during ADP-stimulated ATP synthesis, possibly because electron transport downstream of MQO restores the proton gradient more effectively. When TgCoq4 was downregulated, the malate-driven ADP response became similar to the succinate-driven response, and the membrane potential became more depolarized, likely due to lower UQ levels (**Fig. 6F**, malate -/+ATc).

Taken together, these findings suggest that lower cellular UQ levels after TgCoq4 knockdown limits respiration with both substrates. However, malate-supported electron transport maintains the resting membrane potential despite reduced respiration, while succinate-supported electron transport does not. This difference is consistent with our enzymatic assays showing that TgCoq4 knockdown selectively impairs SDH activity, suggesting that TgCoq4 has an additional role in supporting succinate oxidation beyond its function in UQ biosynthesis.

### Potential formation of a higher-order respiratory organization in *T. gondii*

Proximity labeling of TgCoq4 and TgCoq5 also revealed enrichment of CIII subunits, and MbY2H suggested physical interactions between TgCoq4 and Rieske (**Fig. 2**), so we investigated these potential interactions in parasites. To test this, we generated a double-tag line expressing TgCoq4-3HA and Rieske-3Myc and performed reciprocal co-IPs (**Fig. 7A**). TgCoq4 and Rieske co-immunoprecipitated with each other, supporting the interaction identified by MbY2H and providing evidence for a physical association between the UQ synthesis complex and CIII (**Fig. 7B**). While investigating the association between SDH and the UQ synthesis complex using 2DBN/SDS-PAGE, we noticed that, depending on the amount of protein loaded, the signal extended to greater than 2MDa (**Fig. 7C**). To investigate the potential formation of a larger protein assembly, we tagged the CIII Rieske subunit and the CIV ApiCox30 subunit (both enriched in the TurboID experiment) with 3xHA at their C-termini and validated their mitochondrial localization (**Fig. S7A**). Consistent with previous findings [16], CIII and CIV co-migrate when solubilized with the mild detergent digitonin. Similar to mammalian respiratory supercomplexes [28], this supercomplex is destabilized by dodecyl maltoside (DDM) (**Fig. S7B-C**). Two-dimensional BN/SDS-PAGE revealed a similar high-molecular weight complex (>2MDa) containing CIII and CIV subunits (**Fig. 7C**). Taken together, these data suggest that the UQ synthesis complex, SDH, CIII, and CIV may associate in a higher-order respiratory assembly in *T. gondii* (**Fig. 7D**). These findings place the UQ synthesis-SDH interaction within a broader organization of the parasite respiratory chain.

**Figure 7.**
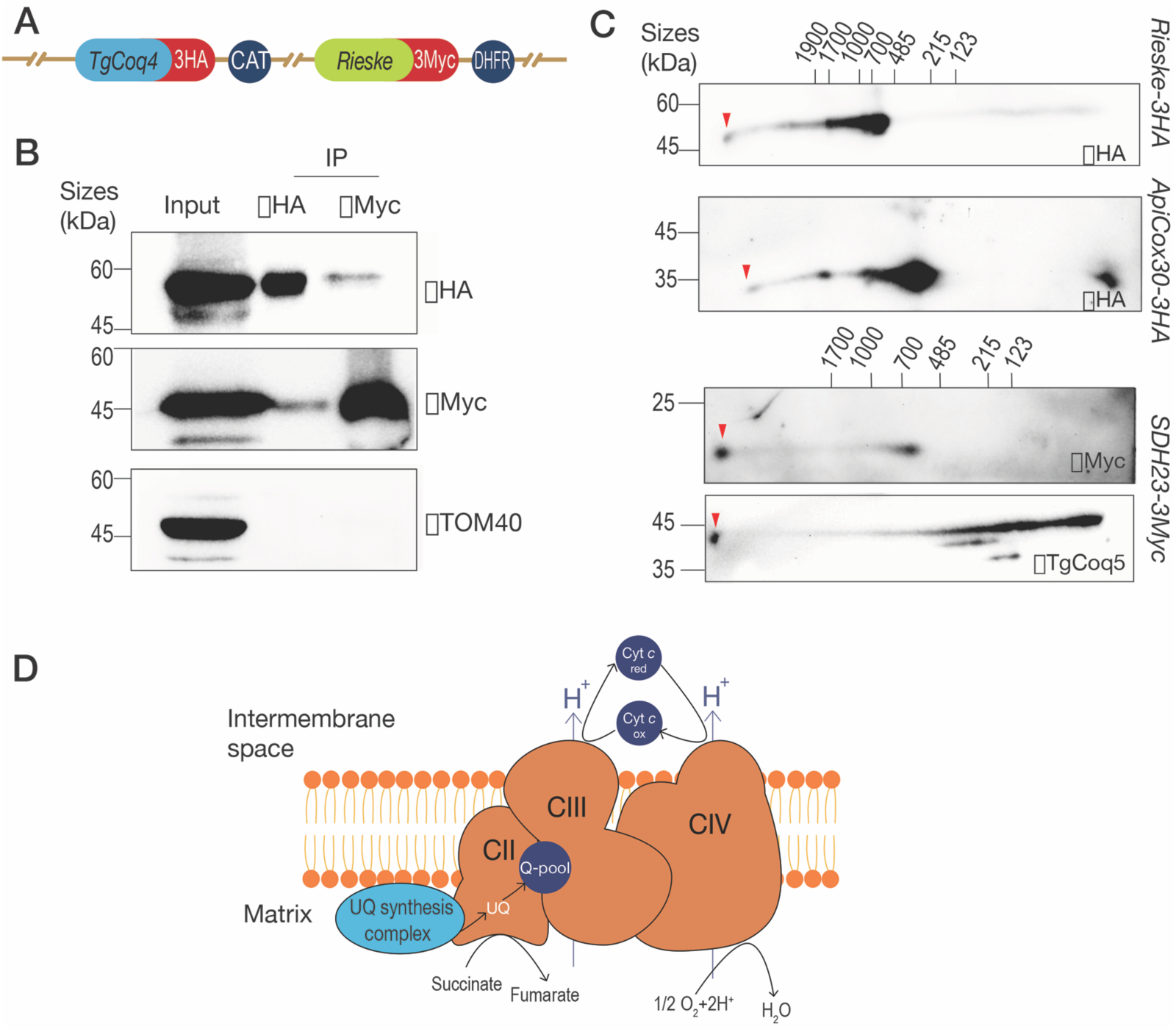
TgCoq4 interacts with the CIII Rieske subunit. (A) Schematic showing the double tagging of *TgCoq4-3HA* and *Rieske-3Myc*. CAT: chloramphenicol acetyltransferase gene used for chloramphenicol resistance; DHFR: dihydrofolate reductase gene used for pyrimethamine resistance. (B) Reciprocal Co-IP using the double-tagged line. aHA was used to pull down TgCoq4-3HA, and aMyc to pull down Rieske-3Myc. TOM40 was used as a negative control. (C) Two-dimensional BN/SDS-PAGE using lysates from *Rieske-3HA*, *ApiCox30-3HA*, and *TgCoq4-3HA*. Membranes were probed with aHA. The red arrowhead indicates the larger protein assembly. (D) Model of the potential larger protein assembly in *T. gondii*. The experiments presented in B and C were repeated at least twice.

## Discussion

Unlike most eukaryotes, apicomplexan parasites lack the canonical mitochondrial CI [12], the major entry point for electrons into the respiratory chain, raising the question of how mitochondrial respiration is organized in these organisms. These parasites have likely evolved multiple adaptations to sustain mitochondrial respiration in the absence of CI. Here, we identify one such adaptation: a parasite-specific organization that physically links ubiquinone synthesis to SDH.

UQ serves an essential role in the mitochondrial inner membrane, transferring electrons from several dehydrogenases to CIII during OXPHOS. The UQH_2_/UQ ratio depends on the balance between electron input and UQH_2_ oxidation by downstream respiratory complexes, as well as ATP demand and oxygen availability [24]. CI is the major entry point for electrons in most eukaryotes [29], but apicomplexan parasites lack CI and use other alternative dehydrogenases to supply electrons to the UQ pool. One challenge faced by parasites lacking CI is how they prevent over-reduction of the UQ pool, particularly when electrons enter through SDH, which can rapidly reduce UQ and may promote ROS accumulation and reduce respiratory efficiency.

Here, we show that *T. gondii* has evolved a previously unrecognized physical and functional microdomain that links the UQ synthesis complex to SDH. This organization may provide SDH with newly synthesized UQ, supporting succinate oxidation and helping to maintain the balance between oxidized and reduced UQ, while revealing a parasite-specific strategy for organizing mitochondrial respiration in the absence of canonical CI.

We show that the UQ synthesis complex physically interacts with SDH subunits in *T. gondii*. TurboID proximity labeling, membrane-based yeast two-hybrid analysis, reciprocal co-immunoprecipitation, and BN-PAGE together support a mitochondrial microdomain that links these two protein complexes. TgCoq4 is essential for UQ synthesis and also contributes to optimal SDH activity, suggesting a direct link between newly synthesized UQ and succinate oxidation. This effect cannot be explained simply by lower UQ levels, since SDH activity remained impaired even in the presence of exogenous UQ_6_. TgCoq5 knockdown did not affect SDH activity, and TgCoq4 knockdown did not impair CIII activity. These findings suggest that TgCoq4 has a specific role in supporting SDH function beyond its role in UQ biosynthesis. The bioenergetic results are consistent with this interpretation. TgCoq4 knockdown decreased respiration with both succinate and malate but reduced the resting mitochondrial membrane potential significantly only with succinate. Malate-supported respiration maintained the resting membrane potential despite lower oxygen consumption. This substrate-specific difference is consistent with the selective impairment of SDH activity following TgCoq4 knockdown and supports a role for the TgCoq4-SDH association in succinate-supported mitochondrial function.

We further show that parasite respiration and ROS production are substrate-dependent, and that succinate oxidation likely limits respiration through over-reduction of the local UQ pool. Importantly, the inability of succinate-driven respiration to reach maximal capacity is alleviated when electrons enter the respiratory chain downstream of SDH or via alternative dehydrogenases, supporting the idea that coupling of UQ biosynthesis to SDH maintains redox balance and prevents UQ-pool bottlenecks. Furthermore, this interaction occurs through parasite-specific SDH subunits, is absent from the mammalian host, and may represent a potential therapeutic target. Interestingly, we also detected interactions between the UQ biosynthesis machinery and CIII, suggesting that UQ synthesis complex may be integrated into a larger respiratory assembly.

In organisms with canonical CI, this complex assembles with CIII and CIV to form a respirasome, which is thought to enhance catalytic activity and stabilize respiratory complexes [2, 6–8, 28]. However, SDH is generally not part of stable respirasomes, although chemical crosslinking has detected weak or transient interactions between SDH and other ETC complexes [9]. In the canonical respirasome, CI transfers electrons from NADH to UQ, reducing it to UQH_2_, which then transfers electrons to CIII which then reduces cytochrome c through the Q cycle. In our model, in *T. gondii*, SDH reduces UQ to UQH_2_, which transfers electrons to CIII for transfer to cytochrome c through the Q cycle. However, since succinate tends to over-reduce the UQ pool, the microdomain between the UQ synthesis complex and SDH might help prevent this by supplying SDH with newly synthesized oxidized UQ and/or by regulating SDH activity. Lack of canonical CI may also be beneficial for these intracellular parasites, as CI is a major site of mitochondrial ROS production [30], and its absence may lower intrinsic ROS production by eliminating one of the major sources of ROS, helping to limit oxidative damage during parasite metabolism. Excessive ROS could diffuse or be converted to hydrogen peroxide that can reach the host cytosol and amplify immune signaling [31–33]. Hence, the lack of canonical CI, together with a specialized microdomain between UQ synthesis and SDH, may represent an evolutionary strategy to sustain respiration while limiting ROS production.

In this work, we assessed the UQ redox balance indirectly by measuring hydrogen peroxide production. Future work should directly measure the UQ redox state. These results raise the possibility that organizing UQ biosynthesis in proximity to SDH or other dehydrogenases may represent a general strategy to buffer the UQ redox state and support respiration when CI function is compromised. This possibility could be explored in models of human CI deficiency or dysfunction.

Together, these findings identify a potential mechanism by which *T. gondii* organizes electron transport without canonical CI, highlighting the UQ synthesis-SDH interaction as a potential therapeutic target.

## Materials and Methods

### Culture

hTERT (human telomerase reverse transcriptase) cells were cultured in Dulbecco’s modified minimal essential medium (DMEM) supplemented with 10% BCS (Gemini Bio-Products, 100-506-500). Cell cultures were maintained at 5% CO_2_ at 37°C. *T. gondii* tachyzoites were cultured in hTERT cells DMEM with 1% BCS and purified as described previously[34].

The C-terminally tagged line mixed populations were cultured in DMEM with 1% BCS with 6.8 µg/mL Chloramphenicol (Sigma, C0378). The conditional knockdown mixed populations were cultured in DMEM with 1% BCS containing 1 µM pyrimethamine (Sigma, P7771). The clonal lines were cultured in DMEM with 1% BCS. For downregulation of the genes, 0.5 µg/mL anhydrotetracycline (ATc) (Cayman Chemical Co Inc, 10009542-500) was added.

### Generation of mutant parasites

For C-terminus tagging, we used CRISPR/Cas9-based genome editing [35] to introduce a 3xHA tag and a chloramphenicol drug resistant cassette into the 3’ end of the gene of interest (GOI) upstream to the stop codon. We used a single guide RNA targeting the 3’ end of the open reading frame into pSAG1-CAS9-U6-sgUPRT [36] using Q5-site directed mutagenesis (NEB, M0491L). The primers used for all mutants are listed in the Supplemental Table 1, and we confirmed the insertion by Sanger sequencing. For each GOI, we amplified a donor DNA sequence encoding 3xHA from pLic-3HA-CAT [37, 38] or 3xMyc from pLIC-3Myc-DHFR (generated in this study) in tandem with 40 bp flanks of the GOI as listed in Supplementary table 1. For generation of TurboID tagged lines, we amplified a donor DNA sequence encoding TurboID-3HA from the 3xHA TurboID pLIC plasmid (modified from Addgene #107169) in tandem with 40 bp flanks of the 3’ end of *TgCoq4* (TGGT1_275980).

For generation of conditional knockdown cell lines, we used a single guide RNA targeting the 5’ open reading frame of the GOI into the pSS013 [35] plasmid using restriction enzymes as previously described [35]. The primers used are listed in Supplementary table 1, and the sequences were confirmed by sequencing. For each GOI, we amplified the donor DNA sequence encoding Tet7SAG4 and a pyrimethamine drug resistant cassette from the pDTS4myc plasmid [37] in tandem with 40 bp flanks of the GOI using primer pairs listed in Supplementary table 1. For each cell line, we transfected the modified CAS9 plasmid and donor DNA by electroporation into *TatiΔku80* or the C-terminus tagged line as mentioned in the results. All primers used in this study are listed in Supplementary Table 1.

### Immunofluorescent microscopy (IFA) and western blots

For IFAs with intracellular tachyzoites, sub-confluent hTERT cells seeded onto coverslips were infected for 20-24 h and the coverslips fixed with 3% paraformaldehyde for 15 min and permeabilized with 0.25% Triton X100 for 10 min. The slides were blocked in PBS pH 8.0 containing 3% bovine serum albumin (BSA) for 30 min prior to incubation with the respective antibody. Primary and secondary antibodies were incubated for 1 h each. Antibodies were diluted in PBS pH 8.0 with 3% BSA. Primary rat αHA (Roche, 11867423001) was used at 1:25, rabbit αTOM40 [39] (gift from the van Dooren lab) was used at 1:20,000, rabbit αVP16 (Santa Cruz biotech, sc-7545) was used at 1:100. Secondary antibodies include GαM 546 (Invitrogen, A-11030), GαRab 488 (Invitrogen, A-11006), and Streptavidin 488 (Invitrogen, S32354) and were used at 1:1,000. Coverslips were mounted on glass slides with DAPI fluoromount-G. Images were taken with a Delta Vision microscope (Olympus, IX-71) with a Photometric Coolsnap camera. Deconvolved images were obtained by using softWoRx deconvolution software.

For western blots, PAGE separated proteins were transferred onto PVDF and blocked in 5% non-fat milk in PBS-T (0.1% Tween-20 in PBS) for 30 min. Incubation with primary antibodies was done overnight at 4°C, and secondary antibodies for 1 h at room temp. Antibodies were diluted in PBS-T. Primary rat αHA (Roche, 11867423001) was used at 1:200, αHA (bioLegend, 901533) was used at 1:1,000, mouse αMyc (Santa Cruz biotech, sc-40) at 1:100, rat αMyc (Invitrogen, MCA1929) at 1:500, rat αSAG1 at 1:1,000 (gift from the Vern Carruthers lab), mouse αTy at 1:1,000 (gift from the Diego Huet lab), and mouse αPDIA3 at 1:5,000 (made in the Moreno lab). Secondary antibodies used were αMouse HRP (Invitrogen, 61-6520), αRat HRP (Invitrogen, 31470), and αRabbit HRP (Invitrogen, 31460) at 1:10,000. Proteins were then imaged with a chemiluminescence using ChemiDoc (Bio-Rad, ChemiDoc^TM^ Imaging system).

### Pulldown of biotinylated proteins

Subcellular fractionation and pulldown assays were performed as described [19]. In brief, *TgCoq4-TID* and *FbxO14-TID* lines were incubated with biotin for 1 h prior to fractionation. Mitochondrial enriched fractions were lysed in RIPA buffer (150 mM NaCl, 0.1 % SDS, 0.5 % sodium deoxycholate and 1% NP-40 in 50 mM HEPES pH 7.5 buffer) supplemented with mini complete protease inhibitor, and soluble protein concentrations were measured using BCA. 200 µg of proteins were used as input and bound to streptavidin magnetic beads overnight at 4°C. The beads were washed with RIPA buffer, 1 M KCl, 0.1 M Na_2_CO_3_, 2 mM urea, and then twice with RIPA buffer.

### Protein identification using LC-MS/MS

Samples were sent to the Proteomics and Metabolomics Facility, Nebraska Center for Biotechnology, University of Nebraska-Lincoln and analyzed for mass spectrometry (MS) as previously described in[40]. Briefly, an aliquot of 37.5 µL of sample was added to 12.5 µL of 4X reducing NuPAGE LDS gel (Thermo Fisher Scientific, Waltham, MA) sample buffer at 5 mM dithiothreitol (DTT) and incubated at 95°C for 10 min. The samples were loaded and run on a Bolt 12% Bis-Tris-Plus gel (Thermo Fisher Scientific) in MES SDS running buffer to clean-up the samples and concentrate the proteins into the top of the gel. The gel was then fixed in methanol:acetic acid:water (40:10:50) and stained with Colloidal Coomassie blue G-250. The gel containing proteins was excised and destained in 50% acetonitrile (ACN), 50 mM ammonium bicarbonate. The proteins were reduced in 100 mM ammonium bicarbonate with DTT at 10 mM. The reducing buffer was removed, and proteins were alkylated with iodoacetamide 10 mM. Proteins were digested with 250 ng of trypsin overnight at 37°C. Peptides were extracted from the gel pieces, dried down, and re-dissolved in 5% acetonitrile, 0.2% formic acid. Each digest was run by nanoLC-MS/MS using a 2 h gradient on a Waters CSH 0.075 mm x 250 mm C18 column (Waters Corp, Milford, MA) feeding into a Thermo Orbitrap Eclipse mass spectrometer.

All MS/MS samples were analyzed using Mascot (Matrix Science, London, UK; version 2.7). Mascot was set up to search the cRAP_20150130.fasta (125 entries); and ToxoDB-59_TgondiiGT1_AnnotatedProteins_20221003 (8,460 sequences) assuming the digestion enzyme trypsin. Mascot was searched with a fragment ion mass tolerance of 0.060 Da and a parent ion tolerance of 10.0 PPM. Carbamidomethyl of cysteine was set as a fixed modification. Deamidated of asparagine and glutamine, oxidation of methionine was specified in Mascot as variable modifications. Scaffold (version Scaffold_4.8.9; Proteome Software Inc., Portland, OR) was used to validate MS/MS-based peptide and protein identifications.

Hits were identified with 95% protein threshold and peptide threshold, and the minimum number of peptides detected was set at 1. Statistical analysis was done using Fisher’s exact test with Benjamini-Hochberg multiple test correction to identify significantly enriched proteins.

### Membrane based yeast two-hybrid (MbY2H) assays

This assay was conducted as described [21, 41]. Truncated cDNA of *TgCoq4* (TGGT1_275980), *TgCoq5* (TGGT1_266850), *TgSDHB* (TGGT1_215280), *TgSDH10* (TGGT1_227290), *TgSDH23* (TGGT1_206480), *TgSDH31* (TGGT1_306650), and *TgRieske* without the sequences encoding the putative mitochondrial targeting signal were amplified from *T. gondii* cDNA using the corresponding primers including SfiI restriction site (see supplementary table 1 for primers used). The amplified cDNA, the MbY2H bait plasmid (pBT3-SUC), and the MbY2H prey plasmid (pPR3N) were digested with SfiI and ligated to generate MbY2H constructs, which were confirmed by sequencing. Generation of the TgCoq4 triple mutant was done using the Q5 site directed mutagenesis kit using primers listed in Supplementary Table 1 and mutations were confirmed via sequencing. The bait and prey plasmids were then co-transformed into the yeast NMY51 strain, by incubation at 30°C for 3 to 4 days. The colonies were used for plate and β-galactosidase assays. Bait and empty prey co-transformants were first cultured on SD-4DO with different concentrations of 3-amino-1,2,4-triazole (3AT), a histidine analog and competitive inhibitor of the *His3* gene product, to screen for bait auto-activation. Optimal 3AT concentrations were determined for each bait construct and used for plate assay as described and β-gal assay as described by the manufacturer’s protocol (ThermoFisher catalog number: 75707). For each co-transformant, 9-14 individual colonies were tested, and statistical significance was determined using two-way ANOVA test.

### Reciprocal immunoprecipitations

For reciprocal co-IPs, IPs were carried out as described previously [19] with modifications. Parasites were lysed with 1% digitonin, 20 mM HEPES pH 7.4, 200 mM NaCl, 2 mM MgCl_2_ supplemented with mini complete protease inhibitor on ice for 1 h, and lysates were clarified by centrifugation at 21,000 x g for 15 min at 4°C. A fraction of the clarified lysates were saved as input, and the rest were bound to αHA magnetic beads (Thermo Fisher, 88836), or αMyc magnetic beads (Sigma-Aldrich, SAE0201) overnight at 4°C. The beads were then washed twice with 0.05% digitonin, 20 mM HEPES pH 7.4, 200 mM NaCl, once with 0.05% digitonin, 20 mM HEPES pH 7.4, 500 mM NaCl, followed by another wash with 0.05% digitonin, 20 mM HEPES pH 7.4, 200 mM NaCl before boiling in sample buffer for western blot analysis.

### Native PAGE assays

Double tagged line parasites were solubilized in lysis buffer containing 2% digitonin, 20 mM HEPES pH 7.2, 150 mM NaCl supplemented with mini complete protease inhibitor. Clarified protein lysates were loaded onto the first dimension BN-PAGE gel and solubilized bovine heart mitochondria [42, 43] was loaded as molecular weight marker for mitochondrial complexes. At the end of the run, the lane containing the bovine heart mitochondria was cut off and stained with Coomassie. The lane containing the solubilized P2 fraction was cut and denatured with 1 X Laemmli buffer containing 100 mM DTT and heated for 20 s in the microwave. After the gel lane was cooled to RT, it was transferred and run in a SDS PAGE gel (second dimension). The protein gel was then transferred onto a PVDF membrane at 25 V, 4°C overnight. The membrane was fixed with acetic acid and de-stained with methanol as previously described [41]. The membrane was then probed with the antibodies mentioned in the western blot section.

### Enzymatic activity assays

SDH and MQO activities were measured as described [17, 20] with modifications. Conditional knockdown mutants were cultured with or without ATc for 3 days before collection. Purified parasites were resuspended in a mitochondrial assay buffer (70 mM Sucrose, 220 mM mannitol, 10 mM KH_2_PO_4_, 5 mM MgCl_2_, 1 mM EGTA, 0.2% BSA, in 2 mM HEPES pH 7.2) with 0.2% digitonin at 10^9^ parasite/mL and incubated at 4°C for 30 min. Permeabilized parasites were then added to an assay buffer containing 2 mM succinate or 10 mM Malate, 25 mM KH_2_PO_4_, 0.002175% DCPIP, 1 µM atovaquone, 2.5 mg/mL BSA, and 10 µM UQ_6_. Absorbance at 600 nm was measured every 2 min for 40 min using a plate reader. Enzymatic activity was calculated using the extinction coefficient of DCPIP of 19.1 mM^-1^cm^-1^. Assay buffer without substrates were used as controls.

CIII activity assay was conducted similarly, except the assay buffer contains 25 mM KH_2_PO_4_, 75 µM oxidized equine heart cytochrome c, 100 µM EDTA, 0.0025% Tween-20, 0.5 mM potassium cyanide, 10 µM decylubiquinol. Absorbance at 550 nm was measured every 15 sec for 30 min. CIII activity was measured by using reduced cytochrome c extinction coefficient of 18.5 mM^-1^cm^-1^ [44]. 2.5 µM antimycin A was added as control. Specific activity was calculated by subtracting activity in the presence of antimycin A. Cytochrome *c* reduction assay using succinate or malate as substrate was carried out similarly, except 2 mM succinate or 10 mM malate was used as substrate. For enzymatic assays, the initial linear rate was estimated using GraphPad Prism segmental linear regression to calculate slope 1 as the initial linear rate.

### Mitochondrial membrane potential and oxygen consumption

*TatiΔKu80* and *iΔTgCoq4-3HA* were used to infect confluent hTert in T75 flasks and cultured in media with or without ATc for two days. Freshly egressed parasites were collected, washed with BAG (116 mM NaCl, 5.4 mM KCl, 0.8 mM MgSO_4_, 50 mM HEPES pH 7.3 with 5.5 mM Glucose) and counted. For mitochondrial oxygen consumption, we used protocols previously described [12, 34, 45, 46].

The mitochondrial oxygen consumption rate was obtained using an Oroboros O2K respirometer (https://www.oroboros.at/). 10^8^ parasite was added to one of the chambers containing intracellular buffer 125 mM sucrose, 65 mM KCl, 1 mM MgCl_2_ and 2.5 mM KH_2_PO_4_ in 10 mM HEPES-KOH buffer pH 7.2) and substrates as indicated in the figure legend (Succinate: 2 mM, malate: 10 mM, TMPD/Asc: 0.2 mM TMPD and 1.5 mM ascorbate). Parasites were permeabilized with 12.5 µM digitonin, and the basal mOCR was measured. Addition of 200 µM ADP would stimulate the mOCR, 1.5 µM of oligomycin was added to obtain the minimal mOCR, and lastly 2 µM (when succinate is used) or 10 µM (when malate or TMPD/Asc is used) FCCP was added to depolarize the mitochondrial membrane potential. Mitochondrial membrane potential was measured at the same time as oxygen consumption using 2.5 µM Safranin *O* as a fluorescent probe. Its fluorescence was measured with Fluo Smart-Module. Data was analyzed using DatLab 8.3.

### Amplex UltraRed hydrogen peroxide production assay

Experiments were conducted as per manufacturer’s protocol (catalog number: A12222). Parasites were collected and permeabilized as described in the enzymatic activity section, and the working solution contains 100 µM Amplex UltraRed, 0.2 U/mL HRP, 24 U/mL SOD. Substrates and inhibitors concentrations used were: 2 mM succinate or 10 mM malate, 2.5 µg/mL oligomycin, 2.5 µM atovaquone, 2.5 µM antimycin A. 2 µM H_2_O_2_ was used as internal positive control and the negative control consisted of the mitochondrial assay buffer. Parasite lysates without substrates were used as baseline. Fluorescence was measured at 530/590 nm every 30 sec for 40 min. The baseline fluorescence increase was subtracted from each condition, and the resulting values were normalized to the no-inhibitor condition.

### Statistical analysis

Experimental data were expressed as the mean with standard error of the mean (SEM) from at least three biological repeats unless indicated otherwise. Statistical analyses were performed using GraphPad PRISM version 10. Student’s t-test, one-way ANOVA, and two-way ANOVA tests were used when appropriate and indicated in the figure legends. Significance was considered when *p*-value was less than 0.05.

## Acknowledgements

We thank the Proteomics & Metabolomics Facility (RRID:SCR_021314) at the Nebraska Center for Biotechnology at the University of Nebraska–Lincoln for the mass spectrometry analysis. The facility and instrumentation are supported by the Nebraska Research Initiative. We thank the CTEGD biomedical microscopy cores for the use of their facilities and equipment. Dr. Muthugapatti K. Kandasamy provided training and support. The *FBXO14-TID* cell line used as control for the TURBO experiments was a gift from Diego Huet’s lab. The Fluo Smart-Module was a gift from Dr. Jarred Call from UGA. The TOM40 antibody was a gift from Giel Van Dooren.

This work was funded by NIH grant AI169846 to S.N.J.M. B.B. was partially funded by AHA predoctoral fellowship 24PRE1192541.

## Author contributions

B.B performed all the experiments, analyzed the data and wrote the manuscript. S.N.J.M. supervised the project and edited the manuscript.

## References

1. Mitchell, P., Chemiosmotic coupling in oxidative and photosynthetic phosphorylation. Biol Rev Camb Philos Soc, 1966. 41(3): p. 445–502.

2. Lobo-Jarne, T. and C. Ugalde, Respiratory chain supercomplexes: Structures, function and biogenesis. Semin Cell Dev Biol, 2018. 76: p. 179–190.

3. Lenaz, G. and M.L. Genova, Structure and organization of mitochondrial respiratory complexes: a new understanding of an old subject. Antioxid Redox Signal, 2010. 12(8): p. 961–1008.

4. Mantle, D., M. Dewsbury, and I.P. Hargreaves, The Ubiquinone-Ubiquinol Redox Cycle and Its Clinical Consequences: An Overview. Int J Mol Sci, 2024. 25(12).

5. Trumpower, B.L., The protonmotive Q cycle. Energy transduction by coupling of proton translocation to electron transfer by the cytochrome bc1 complex. J Biol Chem, 1990. 265(20): p. 11409–12.

6. Kohler, A., A. Barrientos, F. Fontanesi, and M. Ott, The functional significance of mitochondrial respiratory chain supercomplexes. EMBO Rep, 2023. 24(11): p. e57092.

7. Lenaz, G. and M.L. Genova, Structural and functional organization of the mitochondrial respiratory chain: a dynamic super-assembly. Int J Biochem Cell Biol, 2009. 41(10): p. 1750–1772.

8. Javadov, S., S. Jang, X.R. Chapa-Dubocq, Z. Khuchua, and A.K. Camara, Mitochondrial respiratory supercomplexes in mammalian cells: structural versus functional role. J Mol Med (Berl), 2021. 99(1): p. 57–73.

9. Liu, F., P. Lossl, B.M. Rabbitts, R.S. Balaban, and A.J.R. Heck, The interactome of intact mitochondria by cross-linking mass spectrometry provides evidence for coexisting respiratory supercomplexes. Mol Cell Proteomics, 2018. 17(2): p. 216–232.

10. Hill, D.E., S. Chirukandoth, and J.P. Dubey, Biology and epidemiology of Toxoplasma gondii in man and animals. Anim Health Res Rev, 2005. 6(1): p. 41–61.

11. Nishi, M., K. Hu, J.M. Murray, and D.S. Roos, Organellar dynamics during the cell cycle of Toxoplasma gondii. J Cell Sci, 2008. 121(Pt 9): p. 1559–68.

12. Vercesi, A.E., C.O. Rodrigues, S.A. Uyemura, L. Zhong, and S.N. Moreno, Respiration and oxidative phosphorylation in the apicomplexan parasite Toxoplasma gondii. J Biol Chem, 1998. 273(47): p. 31040–7.

13. Lin, S.S., U. Gross, and W. Bohne, Two internal type II NADH dehydrogenases of Toxoplasma gondii are both required for optimal tachyzoite growth. Mol Microbiol, 2011. 82(1): p. 209–21.

14. Maclean, A.E., H.R. Bridges, M.F. Silva, S. Ding, J. Ovciarikova, J. Hirst, and L. Sheiner, Complexome profile of Toxoplasma gondii mitochondria identifies divergent subunits of respiratory chain complexes including new subunits of cytochrome bc1 complex. PLoS Pathog, 2021. 17(3): p. e1009301.

15. Huet, D., E. Rajendran, G.G. van Dooren, and S. Lourido, Identification of cryptic subunits from an apicomplexan ATP synthase. Elife, 2018. 7.

16. MacLean, A.E., S. Shikha, M. Ferreira Silva, M.J. Gramelspacher, A. Nilsen, K.M. Liebman, S. Pou, R.W. Winter, A. Meir, M.K. Riscoe, J.S. Doggett, L. Sheiner, and A. Muhleip, Structure, assembly and inhibition of the Toxoplasma gondii respiratory chain supercomplex. Nat Struct Mol Biol, 2025. 32(8): p. 1424–1433.

17. Zwahlen, S.M., J.A. Hayward, C.S. Maguire, A.R. Qin, and G.G. van Dooren, A myzozoan-specific protein is an essential membrane-anchoring component of the succinate dehydrogenase complex in Toxoplasma parasites. Open Biol, 2024. 14(6): p. 230463.

18. Brandt, U. and B. Trumpower, The protonmotive Q cycle in mitochondria and bacteria. Crit Rev Biochem Mol Biol, 1994. 29(3): p. 165–97.

19. Baierna, B., T. Rahman, S. Latimer, G.J. Basset, and S.N.J. Moreno, Evolutionary remodeling of ubiquinone biosynthesis in Toxoplasma gondii reveals an essential bi-functional monooxygenase. Nat Commun, 2026.

20. Silva, M.F., K. Douglas, S. Sandalli, A.E. Maclean, and L. Sheiner, Functional and biochemical characterization of the Toxoplasma gondii succinate dehydrogenase complex. PLoS Pathog, 2023. 19(12): p. e1011867.

21. Lentze, N. and D. Auerbach, Membrane-based yeast two-hybrid system to detect protein interactions. Curr Protoc Protein Sci, 2008. **Chapter 19**: p. Unit 19 17.

22. Mockli, N. and D. Auerbach, Quantitative beta-galactosidase assay suitable for high-throughput applications in the yeast two-hybrid system. Biotechniques, 2004. 36(5): p. 872–6.

23. Pesta, D. and E. Gnaiger, High-resolution respirometry: OXPHOS protocols for human cells and permeabilized fibers from small biopsies of human muscle. Methods Mol Biol, 2012. 810: p. 25–58.

24. Murphy, M.P. and E.T. Chouchani, Why succinate? Physiological regulation by a mitochondrial coenzyme Q sentinel. Nat Chem Biol, 2022. 18(5): p. 461–469.

25. Crofts, A.R., The Q-cycle - A Personal Perspective. Photosynth Res, 2004. 80(1-3): p. 223–43.

26. Sarewicz, M. and A. Osyczka, Electronic connection between the quinone and cytochrome C redox pools and its role in regulation of mitochondrial electron transport and redox signaling. Physiol Rev, 2015. 95(1): p. 219–43.

27. Stefely, J.A. and D.J. Pagliarini, Biochemistry of Mitochondrial Coenzyme Q Biosynthesis. Trends Biochem Sci, 2017. 42(10): p. 824–843.

28. Lenaz, G., G. Tioli, A.I. Falasca, and M.L. Genova, Complex I function in mitochondrial supercomplexes. Biochim Biophys Acta, 2016. 1857(7): p. 991–1000.

29. Sharma, L.K., J. Lu, and Y. Bai, Mitochondrial respiratory complex I: structure, function and implication in human diseases. Curr Med Chem, 2009. 16(10): p. 1266–77.

30. Scialo, F., A. Sriram, R. Stefanatos, R.V. Spriggs, S.H.Y. Loh, L.M. Martins, and A. Sanz, Mitochondrial complex I derived ROS regulate stress adaptation in Drosophila melanogaster. Redox Biol, 2020. 32: p. 101450.

31. Maldonado, E., D.A. Rojas, S. Morales, V. Miralles, and A. Solari, Dual and Opposite Roles of Reactive Oxygen Species (ROS) in Chagas Disease: Beneficial on the Pathogen and Harmful on the Host. Oxid Med Cell Longev, 2020. 2020: p. 8867701.

32. Zhou, W., J.H. Quan, Y.H. Lee, D.W. Shin, and G.H. Cha, Toxoplasma gondii Proliferation Require Down-Regulation of Host Nox4 Expression via Activation of PI3 Kinase/Akt Signaling Pathway. PLoS One, 2013. 8(6): p. e66306.

33. Juan, C.A., J.M. Perez de la Lastra, F.J. Plou, and E. Perez-Lebena, The Chemistry of Reactive Oxygen Species (ROS) Revisited: Outlining Their Role in Biological Macromolecules (DNA, Lipids and Proteins) and Induced Pathologies. Int J Mol Sci, 2021. 22(9).

34. Sleda, M.A., Z.H. Li, R. Behera, B. Baierna, C. Li, J. Jumpathong, S.R. Malwal, M. Kawamukai, E. Oldfield, and S.N.J. Moreno, The Heptaprenyl Diphosphate Synthase (Coq1) Is the Target of a Lipophilic Bisphosphonate That Protects Mice against Toxoplasma gondii Infection. mBio, 2022. 13(5): p. e0196622.

35. Sidik, S.M., C.G. Hackett, F. Tran, N.J. Westwood, and S. Lourido, Efficient genome engineering of Toxoplasma gondii using CRISPR/Cas9. PLoS One, 2014. 9(6): p. e100450.

36. Shen, B., K. Brown, S. Long, and L.D. Sibley, Development of CRISPR/Cas9 for Efficient Genome Editing in Toxoplasma gondii. Methods Mol Biol, 2017. 1498: p. 79–103.

37. Sheiner, L., J.L. Demerly, N. Poulsen, W.L. Beatty, O. Lucas, M.S. Behnke, M.W. White, and B. Striepen, A systematic screen to discover and analyze apicoplast proteins identifies a conserved and essential protein import factor. PLoS Pathog, 2011. 7(12): p. e1002392.

38. Huynh, M.H. and V.B. Carruthers, Tagging of endogenous genes in a Toxoplasma gondii strain lacking Ku80. Eukaryot Cell, 2009. 8(4): p. 530–9.

39. van Dooren, G.G., L.M. Yeoh, B. Striepen, and G.I. McFadden, The Import of Proteins into the Mitochondrion of Toxoplasma gondii. J Biol Chem, 2016. 291(37): p. 19335–50.

40. Rinkenberger, N., A. Rosenberg, J.B. Radke, J. Bhushan, T. Tomita, L.M. Weiss, and L.D. Sibley, Susceptibility of Toxoplasma gondii to autophagy in human cells relies on multiple interacting parasite loci. mBio, 2024. 15(1): p. e0259523.

41. Huang, G. and R. Docampo, The Mitochondrial Ca(2+) Uniporter Complex (MCUC) of Trypanosoma brucei Is a Hetero-oligomer That Contains Novel Subunits Essential for Ca(2+) Uptake. mBio, 2018. 9(5).

42. Evers, F., A. Cabrera-Orefice, D.M. Elurbe, M. Kea-Te Lindert, S.D. Boltryk, T.S. Voss, M.A. Huynen, U. Brandt, and T.W.A. Kooij, Composition and stage dynamics of mitochondrial complexes in Plasmodium falciparum. Nat Commun, 2021. 12(1): p. 3820.

43. Usey, M.M. and D. Huet, ATP synthase-associated coiled-coil-helix-coiled-coil-helix (CHCH) domain-containing proteins are critical for mitochondrial function in Toxoplasma gondii. mBio, 2023. 14(5): p. e0176923.

44. Spinazzi, M., A. Casarin, V. Pertegato, L. Salviati, and C. Angelini, Assessment of mitochondrial respiratory chain enzymatic activities on tissues and cultured cells. Nat Protoc, 2012. 7(6): p. 1235–46.

45. Bertolini, M.S., M.A. Chiurillo, N. Lander, A.E. Vercesi, and R. Docampo, MICU1 and MICU2 Play an Essential Role in Mitochondrial Ca(2+) Uptake, Growth, and Infectivity of the Human Pathogen Trypanosoma cruzi. mBio, 2019. 10(3).

46. McPhillie, M.J., et al., Potent Tetrahydroquinolone Eliminates Apicomplexan Parasites. Front Cell Infect Microbiol, 2020. 10: p. 203.

