## Supplementary Material for "A respiratory microdomain links ubiquinone biosynthesis to succinate dehydrogenase in an apicomplexan mitochondrion lacking Complex I"

**Supplementary Table 1 Primers used in this study.**

| Cell line/plasmid | Primer | Sequences |
|---|---|---|
| *TgCoq4-TID* | 275980 3' proto F | /5Phos/AAGTTGCACTCTCTGCGCTTCAGTCCG |
| | 275980 3' proto R | /5Phos/AAAACGGACTGAAGCGCAGAGAGTGCA |
| | 275980 HA homo F | TGCCCACCTCGCGCCTCACGTAGCTCGCGACCAAAGTTCTTACCCGTACGACGTCCCGGA |
| | 275980 HA homo R | TCTTCCGGAAAAGTGAAGAGTGTGTCGAAACAAAGAGAGACCCTCGGGGGGGCAAGAATT |
| pPR3N-TgCoq4 | pPR3N Coq4 SfiI F | ATTAACAAGGCCATTACGGCCGAACCTGACGGCGCG |
| | pPR3N Coq SfiI R | AATGATTGGCCGAGGCGGCCTCAAGAACTTTGGTCGCG |
| pPR3N-TgCoq5 | pPR3N Coq5 SfiI F | ATTAACAAGGCCATTACGGCCGCCAGGCAGACAGCT |
| | pPR3N Coq5 SfiI R | AATGATTGGCCGAGGCGGCCCTACAGGTTTCCTAACTTGAACC |
| pBT3-SUC-SDH10 | SUC SDH10 F | ATTAACAAGGCCATTACGGCCGCGCCATCCACCACG |
| | SUC SDH10 R | AATGATTGGCCGAGGCGGCCCCCTTAGGGACTTTCTTCAAAAACGCA |

| | | |
|---|---|---|
| pBT3-SUC-SDH23 | SUC SDH23 F | ATTAACAAGGCCATTACGGCCCAACTCGTCTCGCGTCGATTC |
| | SUC SDH23 R | AATGATTGGCCGAGGCGGCCCCAAGAGAGACGCCGTACATCAGCT |
| pBT3-SUC-SDH31 | SUC SDH31 F | ATTAACAAGGCCATTACGGCCAGCATTTCGTGTACCCGCTACTC |
| | SUC SDH31 R | AATGATTGGCCGAGGCGGCCCCGTAGAAAAACCGATGCAGACCCG |
| pBT3-SUC-Rieske | SUC Rieske F | ATTaacaaggccattacggccttcgtctacttcatgtcggcttcc |
| | SUC Rieske R | AARGATTGGCGGAGGCGGCCCCGCCCAGCTTGACGGTGTG |
| pLIC-3Myc-DHFR | pLIC_F | ttctatgcacttgcaggatgACGCGCCCTGTAGCGGCGCATTA |
| | pLIC_R | TCGGAGATAAGTTTCTGTTCCCTAGGGAATTCCCGTCCTCCACTTCC |
| | Myc_F | GAGGACGGGAATTCCCTAGGGAACAGAAACTTATCTCCGAG |
| | Myc_R | tttacagcctggcgaagcttTTACAGATCCTCTTCGCTGA |
| | DHFR_F | TCAGCGAAGAGGATCTGTAAaagcttcgccaggctgtaaa |
| | DHFR_R | TGCGCCGCTACAGGGCGCGTcatcctgcaagtgcatagaa |
| *SDHB-3Myc* | SDHB 3' proto F | /5Phos/AAGTTGACCTAGGCATGCCTCCGGGG |
| | SDHB 3' proto R | /5Phos/AAAACCCCGGAGGCATGCCTAGGTCA |
| | SDHB myc homo F | GAAGTCGCAAAAGCTCGCGAAGGACCTTGGCCTCGCTGCGGAACAGAAACTTATCTCCGA |

| | | |
|---|---|---|
| | SDHB myc homo R | GTAAACACCCGATACACATGCACACAGGTTTGCGTGTTCCCATCCTGCAAGTGCATAGAA |
| *iΔTgCoq4-3HA/SDHB-Ty* | SDHB Ty homo F | GAAGTCGCAAAAGCTCGCGAAGGACCTTGGCCTCGCTGCGGAGGTGCACACCAACcaggaccccctggacTAAGACGGAACACGCAAACCTGTGTGCATGTGTATCGGGTGTT |
| | SDHB Ty homo R | AACACCCGATACACATGCACACAGGTTTGCGTGTTCCGTCTTAgtccaggggggtcctgGTTGGTGTGCACCTCCGCAGCGAGGCCAAGGTCCTTCGCGAGCTTTTGCGACTTC |
| *SDH23-3Myc* | SDH23 3' proto F | /5Phos/AAGTTGAGCAACCGGGAATTAGACGG |
| | SDH23 3' proto R | /5Phos/AAAACCGTCTAATTCCCGGTTGCTCA |
| *SDH23-3Myc iΔTgSDHB* | SDH23 myc homo F | GCGCCTCCTTCTTGAAGCTGATGTACGGCGTCTCTCTTGAACAGAAACTTATCTCCGA |
| | SDH23 myc homo R | CCCTACATCCTGTCCCCAACGAATGCGTTCGTTTTTCCCcatcctgcaagtgcatagaa |
| *iΔTgSDHB Rieske-3myc* | SDHB 5' proto F | /5Phos/AAGTTGAAGGGTGGAGAGAAGGCTGG |
| | SDHB 5' proto R | /5Phos/AAAACCAGCCTTCTCTCCACCCTTCA |
| | SDHB PI homo F | TCTCCTTTCCCCCTGTAGCCAAAAGTCCTCTCTGTGCTCTaagcttcgccaggctgtaaa |
| | SDHB PI | AGGCGGCGGGCCTCACGGAGCGAAGCGCAGACGACAACATggttgaagacagacgaaagc |

| | | |
|---|---|---|
| | homo R | |
| *Rieske-3myc* *OR* *Rieske-3HA* | 220 3' proto F | /5Phos/AAGTTGAATGCAAAGCGAAAAACGTG |
| | 220 3' proto R | /5Phos/AAAACACGTTTTTCGCTTTGCATTCA |
| | 220 Myc homo F | ACCGACAGTTTTCCTCGATGACCACACCGTCAAGCTGGGCGAACAGAAACTTATCTCCGA |
| | 220 Myc homo R | TGGTGTGTATGTGAACACAGACAAAAGGCAGGTCTTTTCAcatcctgcaagtgcatagaa |
| | 220 HA homo F | ACCGACAGTTTTCCTCGATGACCACACCGTCAAGCTGGGCTACCCGTACGACGTCCCGGA |
| | 220 HA homo R | TGGTGTGTATGTGAACACAGACAAAAGGCAGGTCTTTTCACCCTCGGGGGGCAAGAATT |
| *ApiCox30-3HA* | 2978103' proto F | /5Phos/AAGTTGTGAGGCAGAGAAATCAAACGG |
| | 2978103' proto R | /5Phos/AAAACCGTTTGATTTCTCTGCCTCACA |
| | 2978103' homo F | AGCTGCCTCTGTGACTGGCGGCGAGAAGAAACAGATTGCGTACCCGTACGACGTCCCGGA |
| | 2978103' homo R | TTGCGAGGGTGTTGCAGACACCCCAGAGACCGAAGTATAGCCCTCGGGGGGGCAAGAATT |

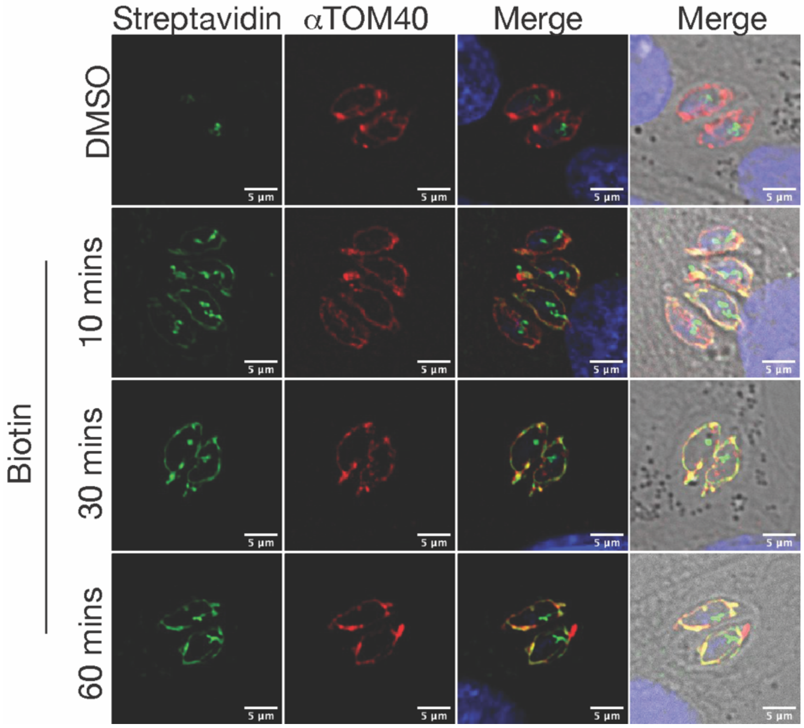


**Supplementary figure 1. *TgCoq4-TID* biotinylates mitochondrial proteins.** The *TgCoq4-TID* line was incubated with or without biotin for 1 h prior to fixation. DMSO was used as the negative control. Streptavidin was used to probe for biotinylated proteins, and TOM40 was used as a mitochondrial marker.

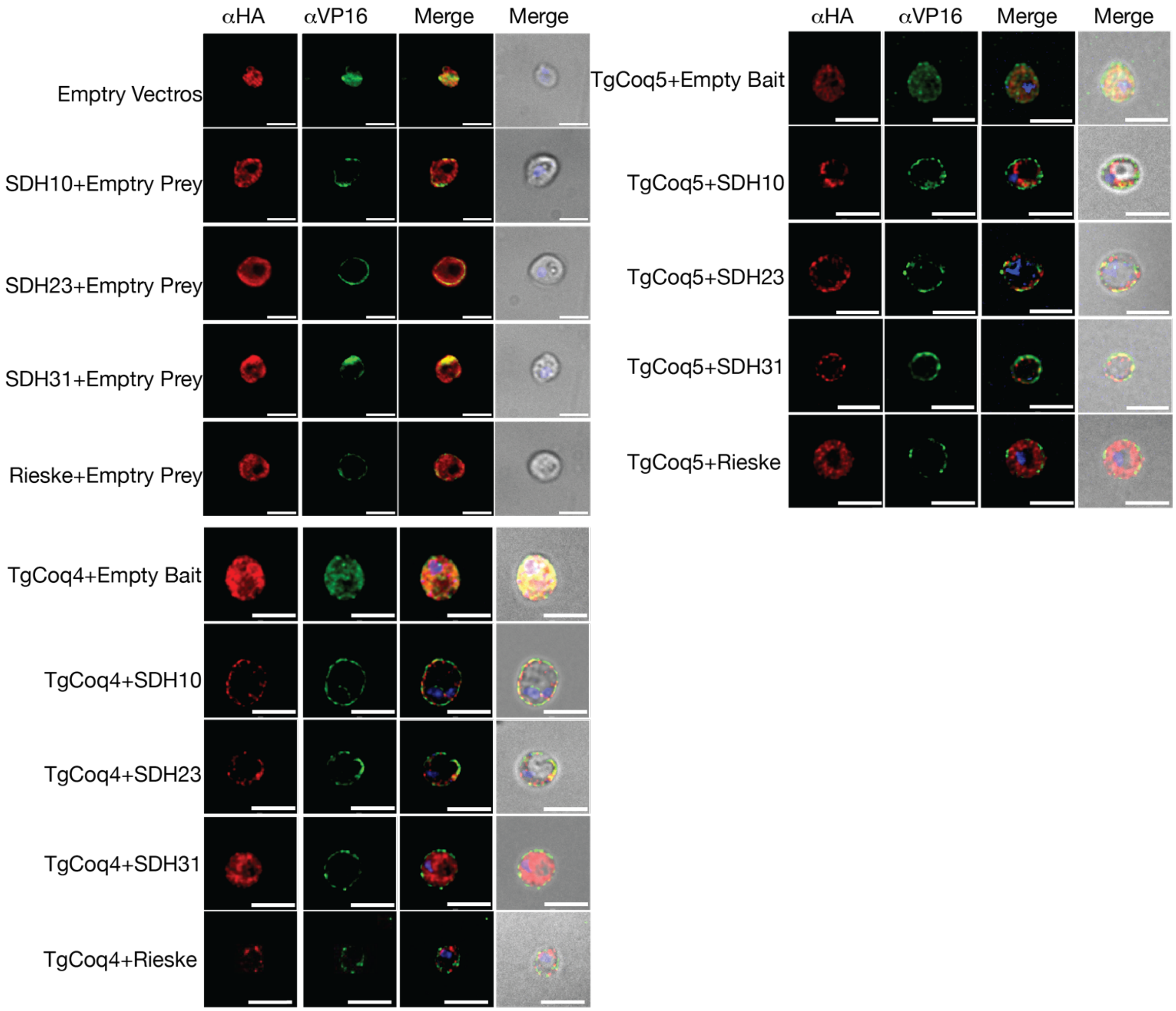


**Supplementary figure 2. MbY2H co-transformants were able to express baits and preys in the correct localizations.** IFA of co-transformants used in **Figure 2**. αHA was used to detect preys, and αVP16 was used to detect baits showing their correct expression at the yeast plasma membrane. Scale bar represents 5 µm.

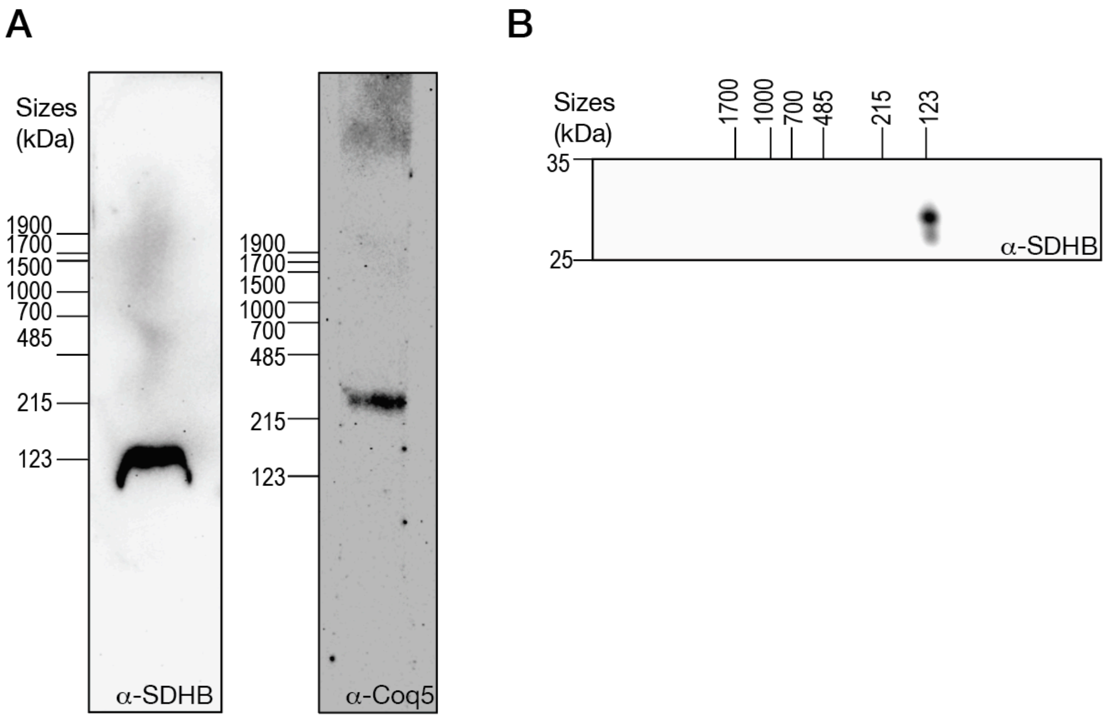


**Supplementary figure 3. SDH and UQ complex migrate separately in HEK cells.** (A) BN-PAGE using HEK cell lysates. SDH was probed using αSDHB, while UQ synthesis complex was detected with αCoq5. (B) Two-dimensional BN/SDS-PAGE using HEK cell lysates.

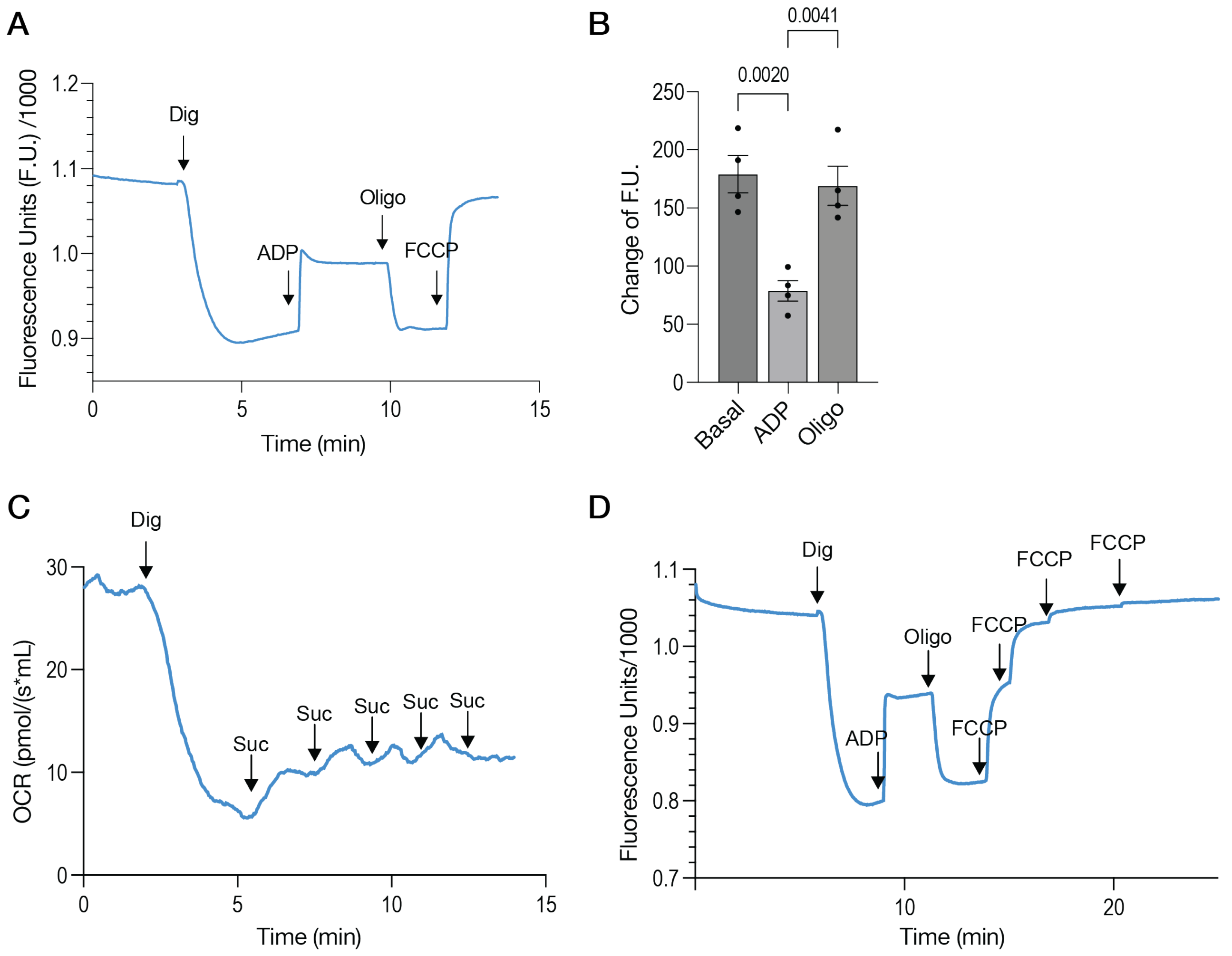


**Supplementary figure 4. FCCP depolarizes the mitochondrial membrane potential when succinate is used as the substrate.** (A) Representative Safranin O fluorescence trace using 2 mM succinate as the substrate. Dig: digitonin; oligo: oligomycin. The concentrations used were the same as those used for the experiments shown in Figure 6. (B) Quantification of the fluorescence (F.U.) changes. Basal: F.U. after FCCP addition minus F.U. before ADP addition, ADP: F.U. after FCCP addition minus F.U. after ADP addition, Oligo: F.U. after the addition of FCCP minus F.U. after oligomycin addition. Data are presented as the mean ± SEM from four biological replicates. A two-way ANOVA was used with statistical significances indicated. (C) OCR trace showing succinate titration. The assay was started without any substrates in the chamber. Dig: digitonin; Suc: succinate. Succinate was added at 0.25 mM per addition. No change was observed after 2 mM. (D) Safranin O fluorescence trace showing FCCP titrations. FCCP was added at 0.5 μM per addition. No change was observed after 2 μM.

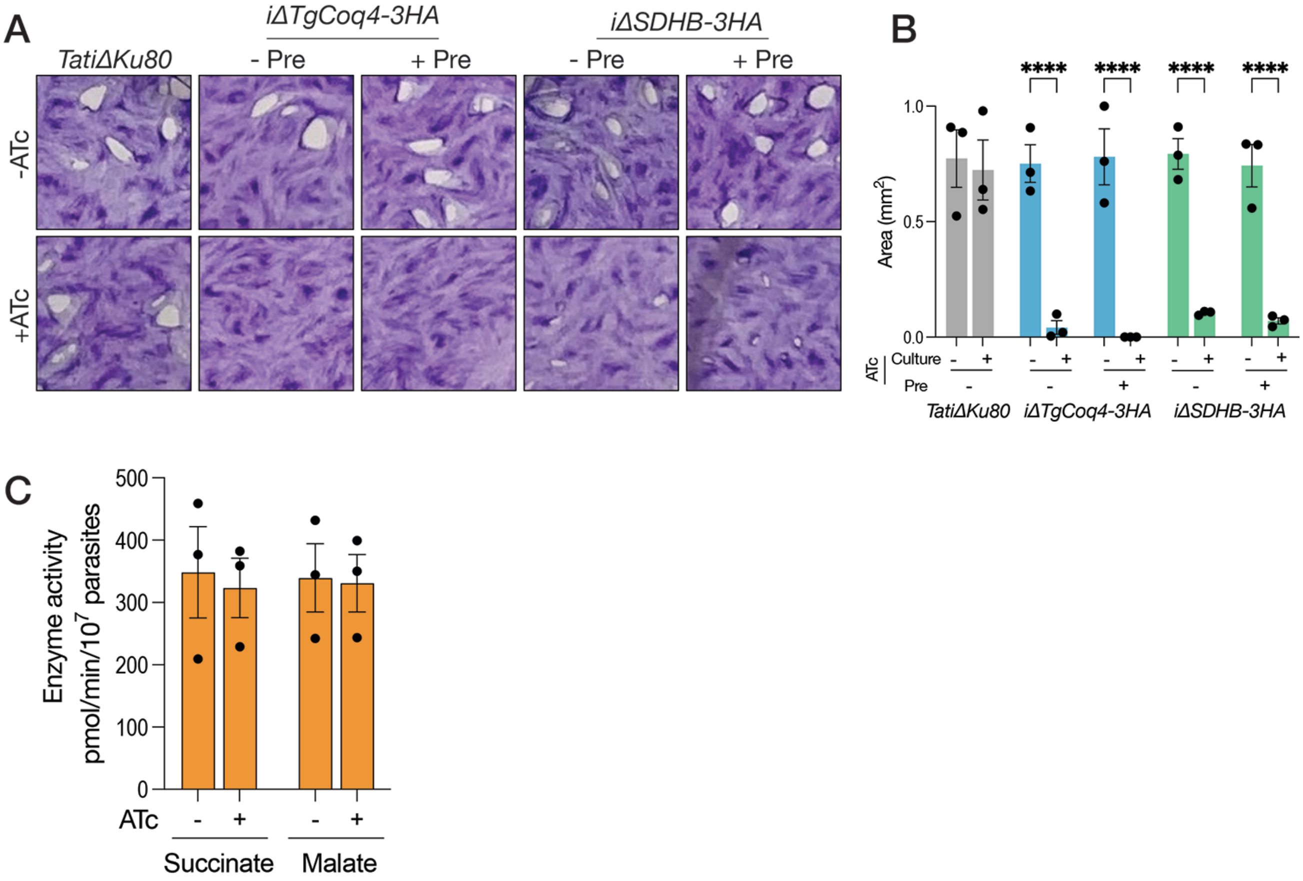


**Supplementary figure 5. TgCoq4 and SDHB are essential for parasite growth, but parasites are still viable after 3 days of ATc treatment.** (A) Representative plaque assay images of *iΔTgCoq4-3HA* and *iΔSDHB-3HA* cell lines either preincubated without or with ATc for 3 days before infection. Parasites were then cultured continuously in media without or with ATc for 7 days before fixation. (B) Quantification of plaque areas from 3 biological replicates. Pre: pre-incubation prior to infection. (C) SDH (using succinate as the substrate) and MQO (using malate as the substrate) activity assays using *iΔTgCoq5* cultured without or with ATc for 3 days. Data are from two biological replicates. For B and C, a two-way ANOVA was used. All data are presented as the mean ± SEM, with each dot representing a biological replicate.

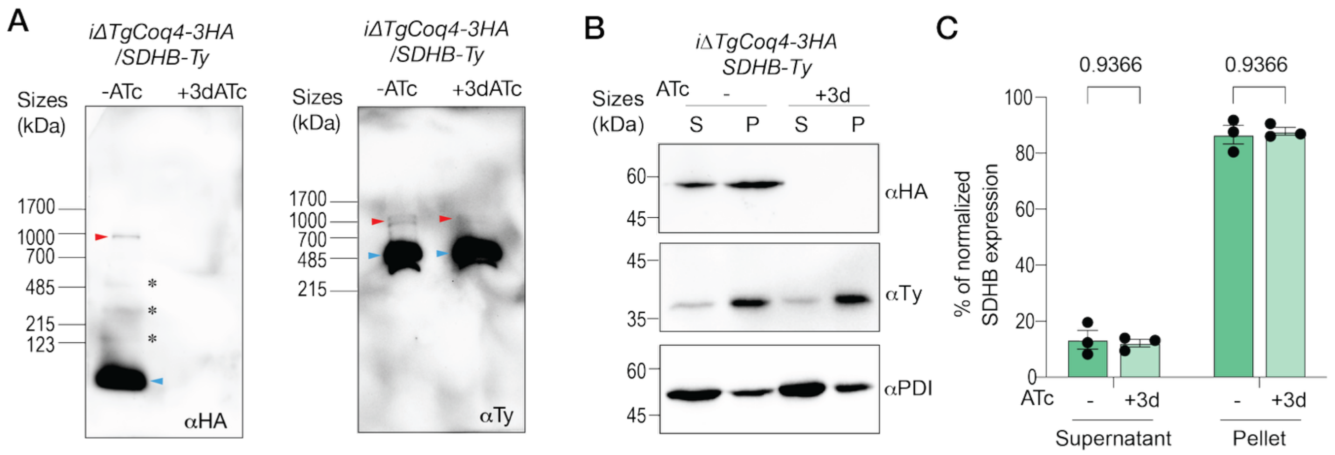


**Supplementary figure 6. TgCoq4 knockdown does not impair SDH assembly or SDHB protein levels.** (A) BN-PAGE using lysates from *iΔTgCoq4-3HA/SDHB-Ty* incubated without or with ATc for 3 days. Left: membrane probed with αHA to detect TgCoq4-3HA; right, membrane was probed with αTy to detect SDHB-Ty. Red arrowhead indicates the 1 MDa supercomplex. Blue arrowhead on the left blot indicates the TgCoq4 monomer, and on the right blot indicates the SDH. Black asterisks indicate UQ synthesis complex assembly intermediates or complex undergoing disassembly during solubilization. (B) Representative western blot using S: soluble fraction and P: pellet after freeze thaw using *iΔTgCoq4-3HA/SDHB-Ty* incubated without or with ATc for 3 days. αHA was used to probe for TgCoq4-3HA, αTy was used to probe for SDHB-Ty, and αPDIA3 was used as loading control. (C) Quantification of the normalized SDHB protein levels (SDHB/PDIA3) shown in B, from three biological replicates. Data are presented as mean ± SEM with each dot representing a biological replicate. A two-way ANOVA was used for statistical analysis.

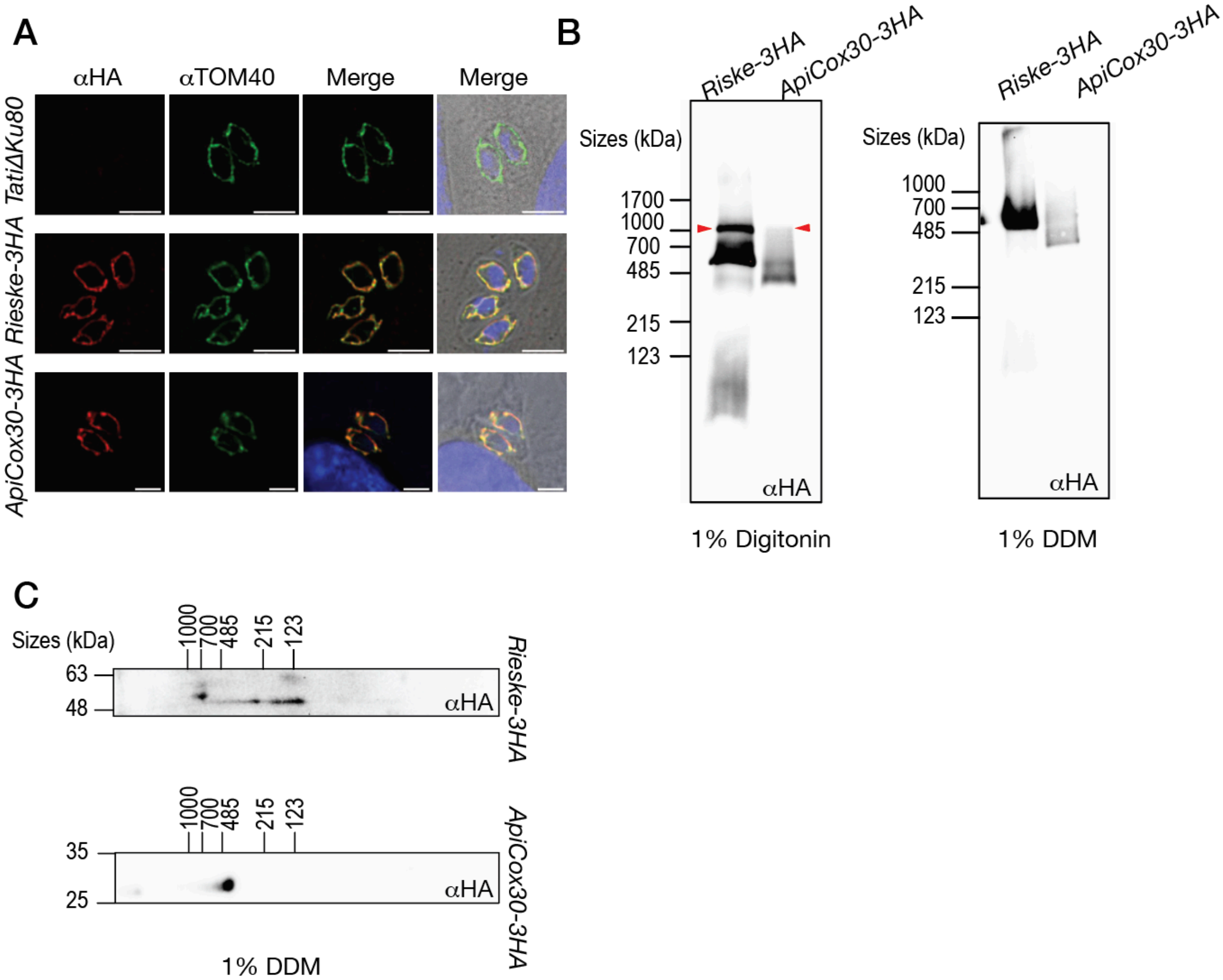


**Supplementary figure 7. CIII and CIV co-migrate when solubilized with digitonin.** (A) IFA validation of the mitochondrial localization of Rieske-3HA and ApiCox30-3HA. TOM40 was used as mitochondrial marker. (B) BN-PAGE using lysates from the indicated parasite lines solubilized with either 1% digitonin or 1% DDM. Red arrowhead indicates the CIII-CIV supercomplex. (C) Two-dimensional BN/SDS-PAGE using parasites solubilized with 1% DDM.
